# Structural basis for ion selectivity in TMEM175 K^+^ channels

**DOI:** 10.1101/480863

**Authors:** Janine D. Brunner, Roman P. Jakob, Tobias Schulze, Yvonne Neldner, Anna Moroni, Gerhard Thiel, Timm Maier, Stephan Schenck

## Abstract

The TMEM175 family constitutes recently discovered K^+^ channels that lack signatures for a P-loop selectivity filter, a hallmark of all known K^+^ channels. This raises the question how selectivity in TMEM175 channels is achieved. Here we report the X-ray structure of a bacterial TMEM175 family member in complex with a novel chaperone built of a nanobody fusion-protein. The structure of the channel in a non-conductive conformation was solved at 2.4 Å and revealed bound K^+^ ions along the channel pore. A hydrated K^+^ ion at the extracellular pore entrance that could be substituted with Cs^+^ and Rb^+^ is coordinated by backbone-oxygens forming a cation-selective filter at the tip of the pore-lining helices. Another K^+^ ion within the pore indicates the passage of dehydrated ions. Unexpectedly, a highly conserved threonine residue deeper in the pore conveys the K^+^ selectivity. The position of this threonine in the non-conductive state suggests major conformational rearrangements of the pore-lining helices for channel opening, possibly involving iris-like motions.

## Introduction

Canonical K^+^ channels constitute a large family of tetrameric membrane proteins that is conserved from bacteria to man mediating the permeability of the membrane for K^+^ ions, crucial to many fundamental processes of the cell. Despite their large structural and functional diversity, all canonical K^+^ channels share the principle of ion selectivity. Their selectivity filter (SF) is built on precisely positioned backbone oxygens in a re-entrant P-loop with the consensus amino acid sequence TVGYG (*1-3*). Numerous parameters were proposed to underlie selectivity, geometrical restraints that disfavor Na^+^ ions because of their size (snug-fit) being the most prominent. In addition, the favored coordination numbers of alkali ions, the carbonyl ligand dynamics and their chemical properties compared to bulk water (dipole) are debated as factors (*1, 4-10*). Recent analyses suggested that desolvated K^+^ ions populate the SF without any intermediate water, and that selectivity arises from the difference in the penalty for dehydration of Na^+^ versus K^+^ (*11-13*). Importantly, selectivity depends on multiple binding sites in the K^+^ SF that transfer ions in file (*12, 14, 15*). Selectivity for ions like Ca^2+^ and Na^+^ is also founded on the P-loop architecture, like in voltage-gated Ca^2+^- and Na^+^-channels (*16-18*) or transient receptor potential (TRP) (*19-21*) channels. However, for these ions high selectivity is also realized in unrelated architectures (*22-24*) whereas K^+^ selectivity, with the exception of the very weakly selective trimeric intracellular cation (TRIC) channels (*25*), is intimately associated with a P-loop architecture.

Recently, members of the transmembrane protein family 175 (TMEM175) have been identified as ion channels, which mediate a major K^+^ permeability in lysosomal membranes and lack a motif related to the P-loop of canonical K^+^ channels (*26*). Given the unequaled prominence of the P-loop SF in K^+^ channels, a very obvious question is how K^+^ conduction is achieved in an unrelated architecture and whether these novel K^+^ channels apply similar principles for selectivity as canonical K^+^ channels. TMEM175 channels are present in animals, eubacteria and archaea. They exhibit selectivities ranging from P_K_/P_Na_ of ~40 in vertebrates to P_K_/P_Na_ of ~2-5 in bacteria and have been described as ‘leak-like’ channels (*26*). The vertebrate proteins are composed of two homologous non-identical repeats, each comprising six transmembrane domains (forming dimers), while the bacterial homologues consist of only one such repeat and form tetramers (*26, 27*). In vertebrates, deletion of the TMEM175 gene leads to increased lysosomal pH under conditions of starvation as well as aberrant autophagosome formation (*26, 28*). The human TMEM175 gene has been linked to Parkinson disease (*28-30*). In prokaryotes, the function of TMEM175 proteins is largely unknown but may be linked to the regulation of the membrane potential (*26*). TMEM175 channels, unlike canonical K^+^ channels, conduct Cs^+^ ions (a hallmark of lysosomal membranes (*31*)) and are neither blocked by Ba^2+^ nor tetraethylammonium or quinine but instead by Zn^2+^ ions. Like other K^+^ channels, they are blocked by 4-aminopyridine (with the exception of bacterial TMEM175 proteins), and conduct Rb^+^ but not Ca^2+^ and N-methyl-D-glucamine (*26, 27*).

## Results

The low similarity to canonical K^+^ channels persuaded us to seek insight into the principles of ion selectivity of TMEM175 channels. We succeeded in obtaining a crystal structure at 2.4 Å resolution, using highly redundant data, of a closed TMEM175 channel from *Marivirga tractuosa* (MtTMEM175). The channel was crystallized in complex with a nanobody (Nb) that was enlarged by fusing Maltose Binding Protein (MBP) to its C-terminus (Nb_51H01-MBPs_) (fig. S1, Table S1, Supplementary Materials and Methods). The overall structure of the MtTMEM175-Nb_51H01-MBPs_-complex is shown in Fig. 1A and fig. S2. We solved the structure by molecular replacement based on this novel Nb-MBP fusion protein that is a promising tool also for other structural biology applications, especially in electron cryo-microscopy. We named this chaperone scaffold ‘macrobody’ (fig. S3, Supplementary Text). Each MtTMEM175 subunit is composed of six transmembrane helices which assemble to form a tetrameric channel, as verified using SEC-MALLS (fig. S4). Nb_51H01-MBPs_ binds extracellular loops distant from the ion conduction path (figs. S2 and S5).

**Fig. 1.**
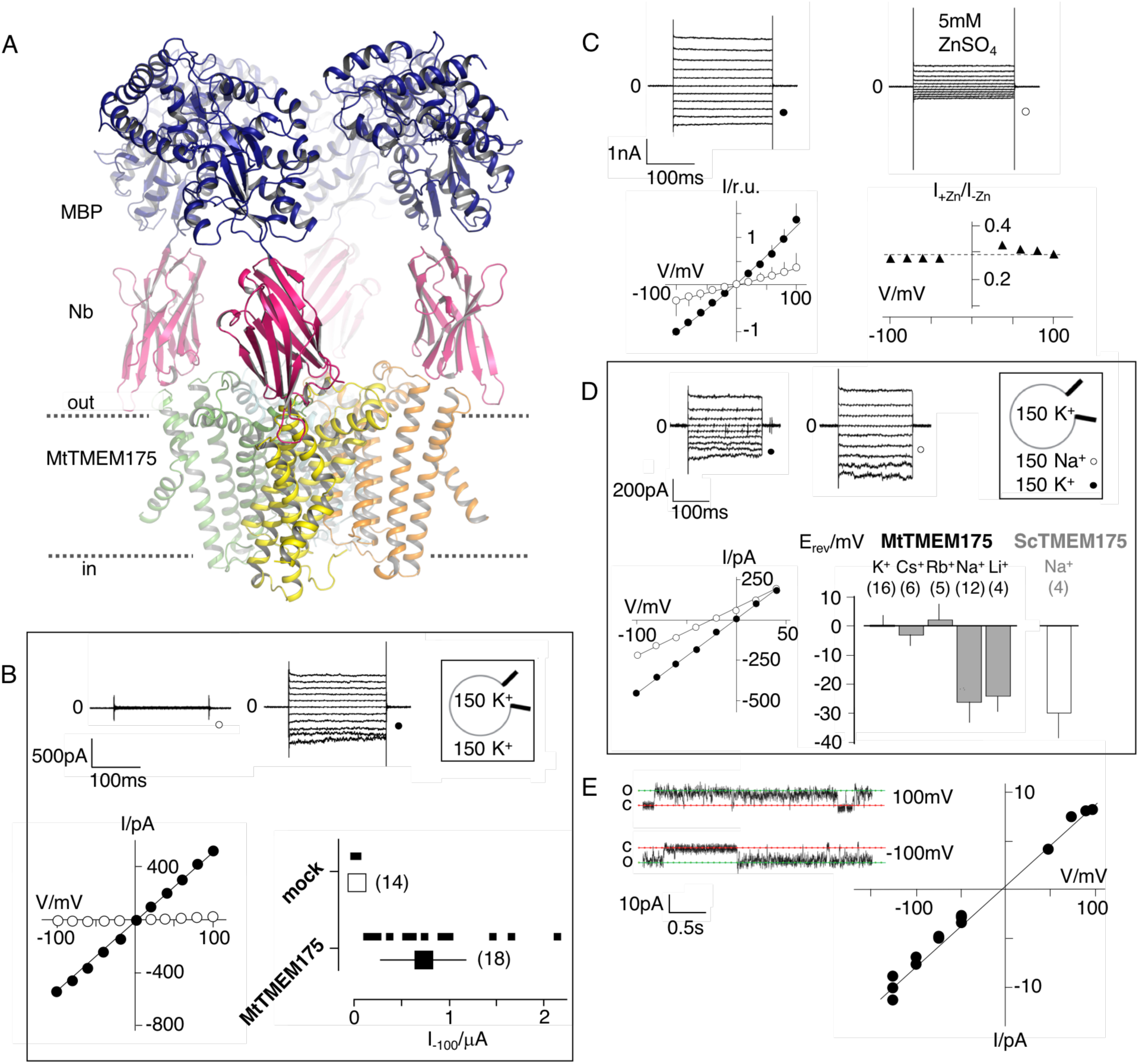
Structure of the MtTMEM175-Nb51H01-MBPs complex and electrophysiological characterization of MtTMEM175. **(A)** Side view of the complex. Approximate membrane boundaries are indicated. **(B)** Current responses to standard voltage pulse protocol (Supplemental Material and Methods) in mock (○) and MtTMEM175 (●) transfected HEK293 cells (upper panel) and corresponding steady state I/V relations (lower left). Plot of currents recorded in same manner at -100 mV for individual cells (small symbols (▪)) and mean ± s.d. (large symbols (□)) (lower right). Number of cells in brackets. **(C)** HEK293 cells expressing MtTMEM175 (top row) before (left) and after (right) adding 5 mM ZnSO_4_ to the bath solution containing 150 mM K^+^. Mean I/V relation (bottom left) of n=4 cells (± s.d.). To compare the effect on different cells the I/V relation was normalized to currents at -100 mV in the absence of blocker (bottom right). **(D)** HEK293 cells expressing MtTMEM175 (top row) before (left) and after (middle) replacing K^+^ (●) with Na^+^ (○) in the external buffer and corresponding I/V relation (bottom left). Same experiments were performed by exchanging K^+^ in external buffer by other cations. The mean reversal voltage (E_rev_) (± s.d., number of cells in brackets) is depicted in right panel for MtTMEM175 and ScTMEM175. **(E)** Exemplary channel fluctuations at ±100 mV measured in cell-attached configuration on HEK293 cells expressing MtTMEM175 (left) and pooled unitary I/V relation of single channel events from measurements in 4 different cells (right) using standard bath and pipette solutions.

For electrophysiological characterization MtTMEM175 was expressed in HEK 293 cells as previously done with homologues from *Streptomyces collinus* and *Chryseobacterium sp.* (ScTMEM175 and CbTMEM175) (*26*). In whole cell patch clamp experiments we recorded non-rectifying, non-inactivating K^+^ currents that showed no signs of voltage-dependence only from transfected cells (Fig. 1B). These currents were blocked by Zn^2+^ ions in a voltage-independent manner (Fig. 1C). Similar to ScTMEM175 and CbTMEM175, MtTMEM175 has also a low selectivity for K^+^ (P_K_/P_Na_ ~2-3). It conducts Cs^+^ and Rb^+^ with a similar efficiency as K^+^, and to lesser extent, similar to Na^+^, also Li^+^ (Fig. 1D and fig. S6). We obtained a few single channel recordings from MtTMEM175-transfected cells that revealed a unit conductance of ~70 pS and showed channel flickering (Fig. 1E). We do not have definitive proof that these currents originate from MtTMEM175, however several arguments support this view. First, we recorded them only in transfected cells, which exhibited typical MtTMEM175 macroscopic currents after breaking into the whole cell configuration. Second, like the macroscopic MtTMEM175 current also the single channel I/V relation reverses around 0 mV (Fig. 1E). Finally, the unitary conductance of the channels at -100mV is in the range of the conductance of mock transfected HEK293 cells in whole cell mode (Fig. 1B), making it unlikely that the currents originate from endogenous channels. The presence of gating events is inconsistent with the definition of a leak channel and has significance for the interpretation of the structure.

Recently, the crystal structure of another bacterial TMEM175 channel was solved at 3.3 Å (CmTMEM175, PDB accession 5VRE), but did not reveal bound ions, even in crystals soaked with heavier monovalent and divalent ions (*27*). In contrast, the structure of MtTMEM175 revealed two densities attributable to K^+^ ions, termed 1K^+^ and 2K^+^ (Fig. 2A, Materials and Methods), supported by data collection at higher wavelengths of 2.02460 Å (fig. S7). One K^+^ ion (1K^+^, occupancy ~1) is located in a short SF at the extracellular pore entrance. It is bound by eight water molecules in an anti-prismatic geometry (fig. S8), which are in turn coordinated by backbone oxygens of Leu^42^, Ser^43^ and Ser^44^ (Fig. 2B). The respective backbone oxygens of these residues are 12, 13.1 and 14.9 Å apart (Fig. 2C). Except for the conserved Leu^42^, no obvious motif for this region is apparent (fig. S9A-B). The second K^+^ ion (2K^+^, occupancy ~0.5) is dehydrated and trapped between the layers of Leu^35^ and Thr^27^ (Fig. 2A), implying that ions pass the channel without hydration shell, and further the existence of a conformation with a wider pore. By soaking the crystals with Cs^+^ and Rb^+^ we detected anomalous density for both ions at the position of 1K^+^ (Fig. 2D) providing additional evidence for selectivity towards monovalent cations with similar properties as K^+^. No significant anomalous signal for Cs^+^ or Rb^+^ was found at the position of 2K^+^, indicating that the channel would have to open for exchange of ions. Additional density in the 2F_o_-F_c_ map on the extracellular side was attributed to a maltose moiety from a detergent molecule. We tested for potential influence of maltose on the conductance by electrophysiology, but could not detect any effects (fig. S10). In comparison with the SF of canonical K^+^ channels, TMEM175 channels comprise a SF of very different architecture. Despite its simplicity, it does however recapitulate a number of central elements seen in K^+^ coordination. Like in canonical K^+^ SFs backbone oxygens are also mediators of coordination in MtTMEM175, but only one ion, surrounded by eight water molecules, resides in the wide filter. This hydrated K^+^ ion is reminiscent of the one in the vestibule of a high-resolution structure of KcsA in close proximity to the SF entrance (*4*). In comparison, the two planes in the K^+^/ water complex in MtTMEM175 are skewed, due to interactions with the surrounding backbone oxygens (fig. S8) (*32*). The eightfold coordination of K^+^ ions in square antiprism geometry (Na^+^ is typically 5-6-fold coordinated) is also seen inside the canonical SF, where it is mediated by backbone oxygens (*33, 34*). MtTMEM175 crystallized in the presence of equimolar amounts of Na^+^ and K^+^ which apparently did not interfere with K^+^ coordination at 1K^+^. Generally, reduction of sequential ion binding sites attenuates the K^+^ selectivity in the canonical K^+^ SF, in both, naturally occurring or mutated SFs whereas introducing additional binding sites can increase selectivity (*10, 35-37*). MtTMEM175 is weakly selective for K^+^ (Fig.1D) which appears conclusive given that only one binding site is present in the SF. In addition the ions do not directly couple to the backbone, which could also contribute to the low selectivity. With its simplicity, the SF of MtTMEM175 is distantly reminding of the short SF of several non-selective TRP channels (*20,38*) but is unique in coordinating a single hydrated K^+^ ion. Further the lack of a vestibule in proximity to the SF is characteristic for TMEM175 channels. Apart from the SF the negative electrostatic potential in the pore lumen is another property promoting cation permeation (fig. S11) (*27*).

**Fig. 2.**
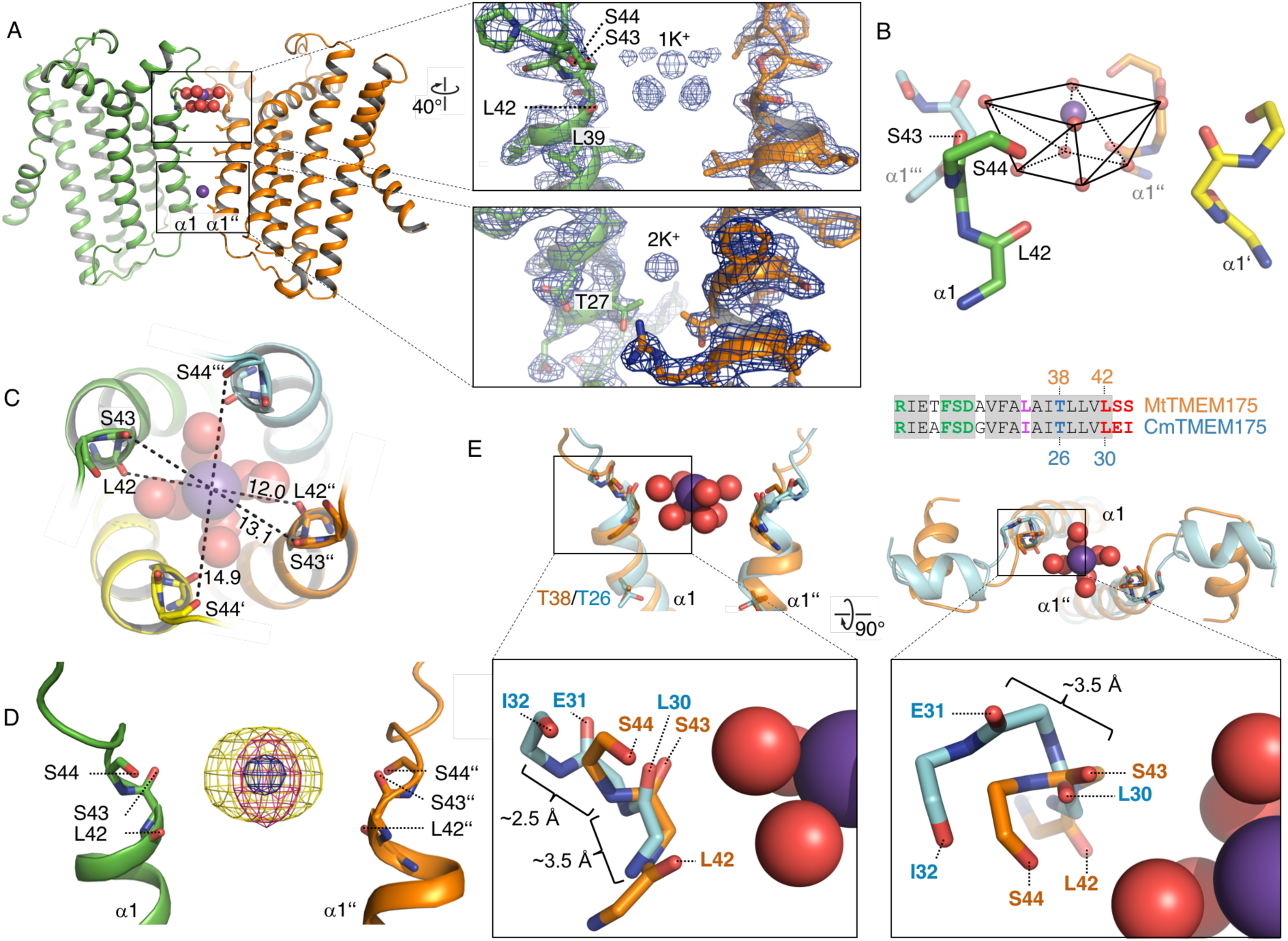
Ion coordination in MtTMEM175. **(A)** Side view (left) and close-up views (right) of the SF with a hydrated K^+^ ion (top) and a trapped K^+^ ion within the pore (bottom). The 2F_o_-F_c_ electron density is depicted as blue mesh (at 2.4 Å, contoured at 1.8 σ, sharpened with b=-25). Two subunits are omitted for clarity. K^+^ ions and water molecules are displayed as purple and red spheres, respectively. **(B)** Interaction of the backbone oxygens of Leu^42^, Ser^43^ and Ser^44^ with the hydrated K^+^ ion and coordination of the K^+^ ion by water molecules in square-antiprismatic geometry (black lines). Side chains are omitted and the size of the spheres is reduced for clarity. **(C)** Top view of the SF. Distances between opposing backbone oxygens of Leu^42^, Ser^43^ and Ser^44^ are indicated in Å. **(D)** Substitution of K^+^ in the SF with Cs^+^ and Rb^+^. The 2F_o_-F_c_ electron density (as in **(A)**, blue mesh) marks the position of the K^+^ ion. Anomalous difference electron densities of Cs^+^ (at 3.8 Å, contoured at 7 σ) and Rb^+^ (at 3.6 Å, contoured at 7 σ, blurred with b=125) are shown in yellow and pink, respectively. **(E)** Superposition of the extracellular ends of helix 1 in MtTMEM175 and CmTMEM175 (5VRE) with approximate deviations indicated in Å (side and top view of the SF). Thr^38^ and Leu^35^ (MtTMEM175, orange) were aligned with Thr^26^ and Ile^23^ (CmTMEM175, cyan). Only main chain atoms are shown. The hydrated K^+^ ion from the MtTMEM175 structure is shown. A sequence alignment of helix 1 is shown for orientation (coloring as in fig. S9).

What is the reason for the absence of ions in the CmTMEM175 structure (*27*)? Intriguingly, the tip of helix 1 in the structure of CmTMEM175 is substantially deviating from the one in our structure, explaining the lack of coordinated ions (Fig. 2E). We found that the extracellular end of helix 1 in the CmTMEM175 structure contains a 3_10_-helix over a stretch of 3-4 residues and thus extends further than helix 1 in MtTMEM175 (fig. S12A-B). 3_10_-helices are associated with transition states and hint to dynamic regions within a helix (*39*), but whether this 3_10_-stretch is an intermediate conformation is unclear. It is worth to note, that the short helix between the helices 1 and 2 of CmTMEM175 is involved in major crystal contacts, which may have caused a displacement of the SF in the preceding loop and formation of the 3_10_-helix stretch (fig. S12C-D).

MtTMEM175 shows a network of hydrogen bonds in proximity to the intracellular pore entrance similar to CmTMEM175 (*27*) (fig. S13, Supplementary Text). Different from CmTMEM175, Arg^24^ is interacting with His^77^ and Asp^30^ of the adjacent subunit, thereby connecting neighboring subunits (fig. S14A). All of these residues are highly conserved (fig. S9). Gel filtration profiles of Arg^24^ mutant proteins support a role in tetramer assembly (fig. S15). In CmTMEM175 this interaction is not observed (fig. S14B), presumably due to protonation of the conserved histidine (CmTMEM175 crystallized at pH 4.6), thereby repelling the arginine.

In the structure of MtTMEM175 Leu^35^ is occluding the pore, similar to the corresponding Ile^23^ in CmTMEM175, to such extent that K^+^ ions cannot pass (Fig. 3A). These bulky residues could thus constitute hydrophobic gates. From single channel recordings and the lack of exchangeability of 2K^+^ with Cs^+^ or Rb^+^ we have indications for open and closed conformations in support of a gate in TMEM175 channels. Opening of the channel could result from displacement of hydrophobic side chains from the pore center, probably by a helix-rotation as seen in the NaK channel (*18*) or in TRPV6 (*21*). Movements in helix 1 would also affect the position of other residues, among them a highly conserved threonine. Threonine^38^ is interspersed between Leu^35^ and Leu^39^ and one of the few polar residues in the pore-lining helix but not facing the pore center. It is participating in a bifurcated hydrogen bond with the main-chain oxygen of Ala^34^ (Fig. 3B). Following this line of thoughts, we reasoned that K^+^ ions could interact with the side chain of Thr^38^ in a conductive conformation of MtTMEM175 and mutated Thr^38^ to alanine (fig. S16A-B). When analyzed in whole cell patch clamp in HEK293 cells the T38A mutant of MtTMEM175 was no longer selective for K^+^ ions, as exchanging K^+^ in the bath solution for Na^+^ caused only a minor shift of the reversal potential by -5.8 ± 3 mV (n=4) (Fig. 3C). For comparison, the WT protein responds to a replacement of K^+^ for Na^+^ with a shift of -27 ± 8 mV (Fig. 1D). The data in Fig. 3C further show that the mutant channel exhibits an anomalous increase in conductance after replacement of K^+^ for Na^+^. The slope conductance between ±40 mV increased in Na^+^ by about 30% (n=4). We thus conclude that Thr^38^ plays a pivotal role for K^+^ selectivity and conductance in TMEM175 channels, reflected also in its high degree of conservation (Fig. S9A-C). Notably, the side chain of a conserved threonine is also essential for the coordination of K^+^ ions at the S4 position in the selectivity filter of canonical K^+^ channels. Hence, not only carbonyl ligands, but also the threonine side chain is suited to coordinate K^+^ ions with impact on selectivity and conductance (*14, 32, 40, 41*). We found no obvious differences in the crystal structure between this mutant and the WT channel in the closed conformation (fig. S16C, Table S2). Despite the clear signature of an eightfold coordinated K^+^ in the extracellular SF from the MtTMEM175 structure, Thr^38^ plays a prominent, if not determining role for selectivity. The extracellular SF could thus serve to attract monovalent cations in general, also with respect to its primordial architecture. However, the function of this extracellular SF/ion binding site might only become fully appreciable upon insight into the conductive conformation which might involve rearrangements at both ends of helix 1. Although we found Zn^2+^ ions by soaking into crystals in the plane of Thr^38^ (fig. S17A), the sensitivity of the channel to Zn^2+^ was not compromised in the T38A mutant. In four cells tested, 5 mM Zn^2+^ blocked the current by 70%±8% and 66%±9% in the WT and mutant, respectively (fig. S17B). Since the Zn^2+^ block is voltage independent (Fig. 1C) this result can be expected, as ion-mediated block inside a pore is commonly voltage dependent (*42*). The location of the Zn^2+^ ion in the structure is probably a result of the attracting electrostatic environment (fig. S11) and the small size of Zn^2+^ and not directly related to the Zn^2+^-block, the nature of which remains obscure for now.

**Fig. 3.**
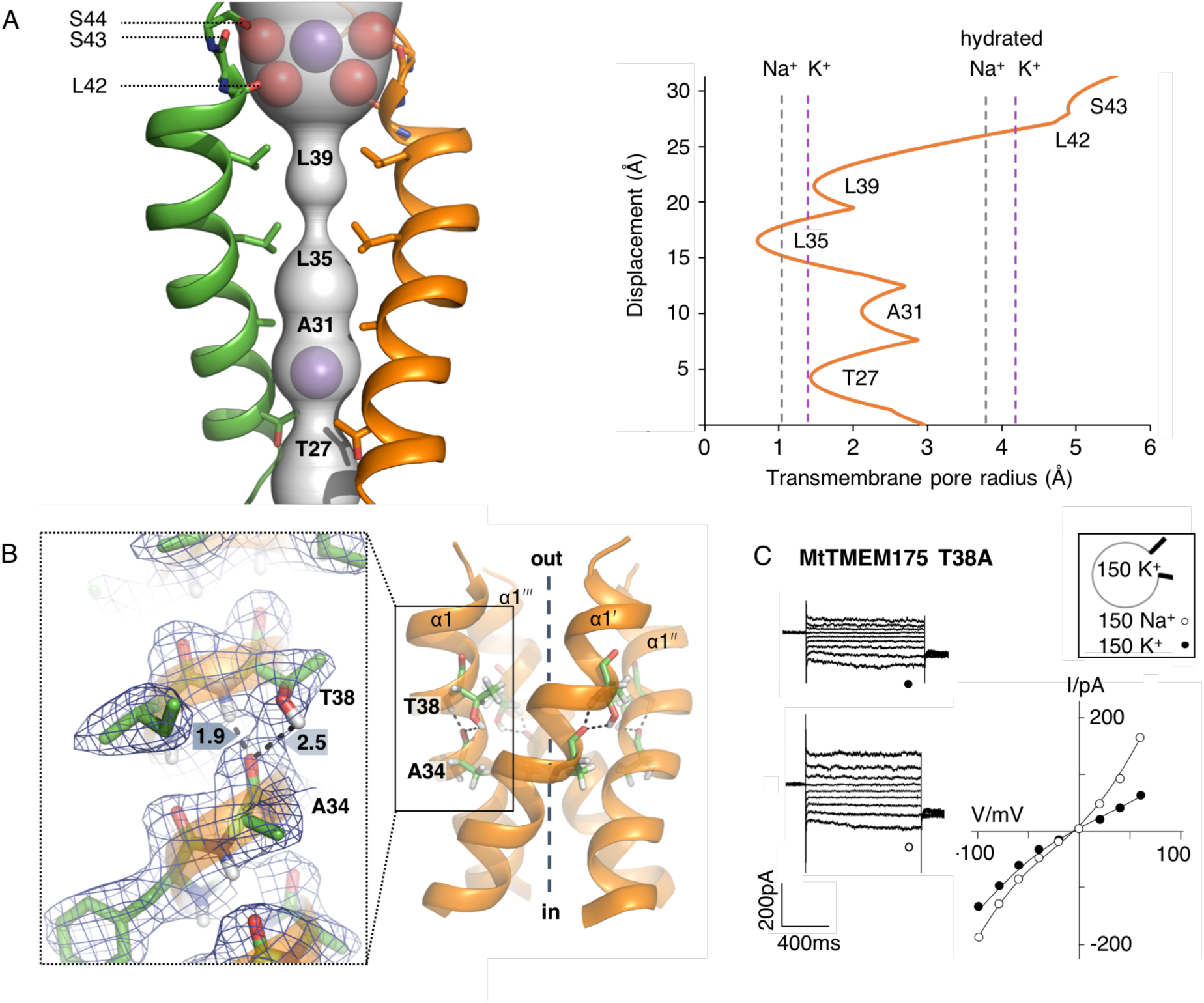
Pore analysis and K^+^ selectivity. **(A)** HOLE analysis. Left: The ion conduction pathway is illustrated as grey surface and pore-lining residues are displayed. K^+^ ions and water molecules are shown as purple and red spheres, respectively. Right: The pore radius along the central axis is shown in Å. Dashed lines indicate the radii of K^+^ and Na^+^ ions with and without hydration shell. **(B)** Position of the highly conserved Thr^38^ in the tetramer. View from the side on the pore-forming helices (right) and close-up view of the bifurcated hydrogen bond between Thr^38^ and Ala^34^ (left). The 2F_o_-F_c_ density (contoured at 1.8 σ after sharpening with b=-25, blue) is displayed. Distances between carbonyl oxygens and hydrogens are given in Å. **(C)** Currents in HEK293 cells expressing the MtTMEM175 T38A mutant before and after exchanging external K^+^ (●) with Na^+^ (○) (left) with corresponding I/V relation (right). The reversal potential is at -5.8 ± 3 mV (n=4).

In contrast to our finding, recent work proposed Ile^23^ in CmTMEM175 to underlie the K^+^ selectivity as part of a hydrophobic selectivity filter. Mutation of Ile^23^ or corresponding residues in hTMEM175 to small or hydrophilic residues resulted in a loss of K^+^ selectivity (*27*). Due to the 3_10_-helix in the CmTMEM175 structure Leu^30^ is facing the pore (*27*), evoking the impression of a triad of pore-lining hydrophobic residues reminiscent of bestrophins, in contrast to the respective residue in the MtTMEM175 structure (Fig. S18). In bestrophins, mutation of the three layers of bulky residues along the ion path to alanine resulted in an open channel, without the requirement for activation (*43, 44*). This supported a function of these residues as a gate instead of contributing to selectivity, contrasting previous interpretations (*45, 46*). Similar observations were made in TWIK-1 where mutation of pore-lining leucine residues to asparagines resulted in a hydrated pore and strongly increased K^+^ currents by interference with the original “vapor-lock” function of residues at these positions (*47*). Also, mutation of a gate built from phenylalanine in the NaK channel strongly increased flux (*18*). If Ile^23^ of CmTMEM175 acts as a gate to keep the channel closed, a role that we suggest for the corresponding Leu^35^ in MtTMEM175, an exchange for asparagine could result in a permanently open channel. Recent simulations of TMEM175 and bestrophins provide an explanation how mutating hydrophobic gates to small and polar residues can turn entirely de-wetted pores into wetted pores that consequently have impact on the closure and conduction of channels (*44, 48*). Recent structures of open states of chicken bestrophin, as determined by cryo-EM, provided evidence for the role of pore-lining hydrophobic residues as gates, convincingly explaining the above-mentioned observations (*49*). Although not purposely intended, mutation of Ile^23^ in CmTMEM175 and equivalent residues in hTMEM175 (*27*) demonstrates that the extracellular SF does not significantly contribute to K^+^ selectivity in agreement with our finding on the T38A mutation in MtTMEM175.

From our analysis we divide the pore of TMEM175 channels into functional layers, markedly different from a previous interpretation (Fig. 4A) (*27*). The MtTMEM175 ion path is thus build from a SF for monovalent cations at the pore tip, a major gate at the position of Leu^35^ and, rather unusual, by an interspersed layer that tunes the selectivity towards K^+^, the highly conserved Thr^38^. The findings regarding Thr^38^ suggest a rotation of helix 1 for channel opening. This view is further fueled by the fact that the most conserved residues along the pore in helix 1 are not facing the pore lumen, but instead the adjacent helix (fig. S19). When viewed from the intracellular side, these small residues (Asp^30^, Ala^34^ and Thr^38^) spiral along the side of the helix and could face the center upon clockwise rotation of helix 1 by a few degrees (Fig. 4B). Such an iris-like motion would simultaneously translocate the side chain of Thr^38^ into the pore and displace the side chain of Leu^35^. A similar scenario for channel opening has recently been reported for bestrophins (*49*) and is conceivable in TMEM175 proteins. Definite conclusions though have to await evidence from an open channel structure and further functional experiments, which should also investigate potential triggers for opening and closing of the channel. Interestingly, the structure of CmTMEM175 shows some aspects of a different state, probably linked to protonation of a conserved histidine (figs. S13-S14 and Supplementary Text).

**Fig. 4.**
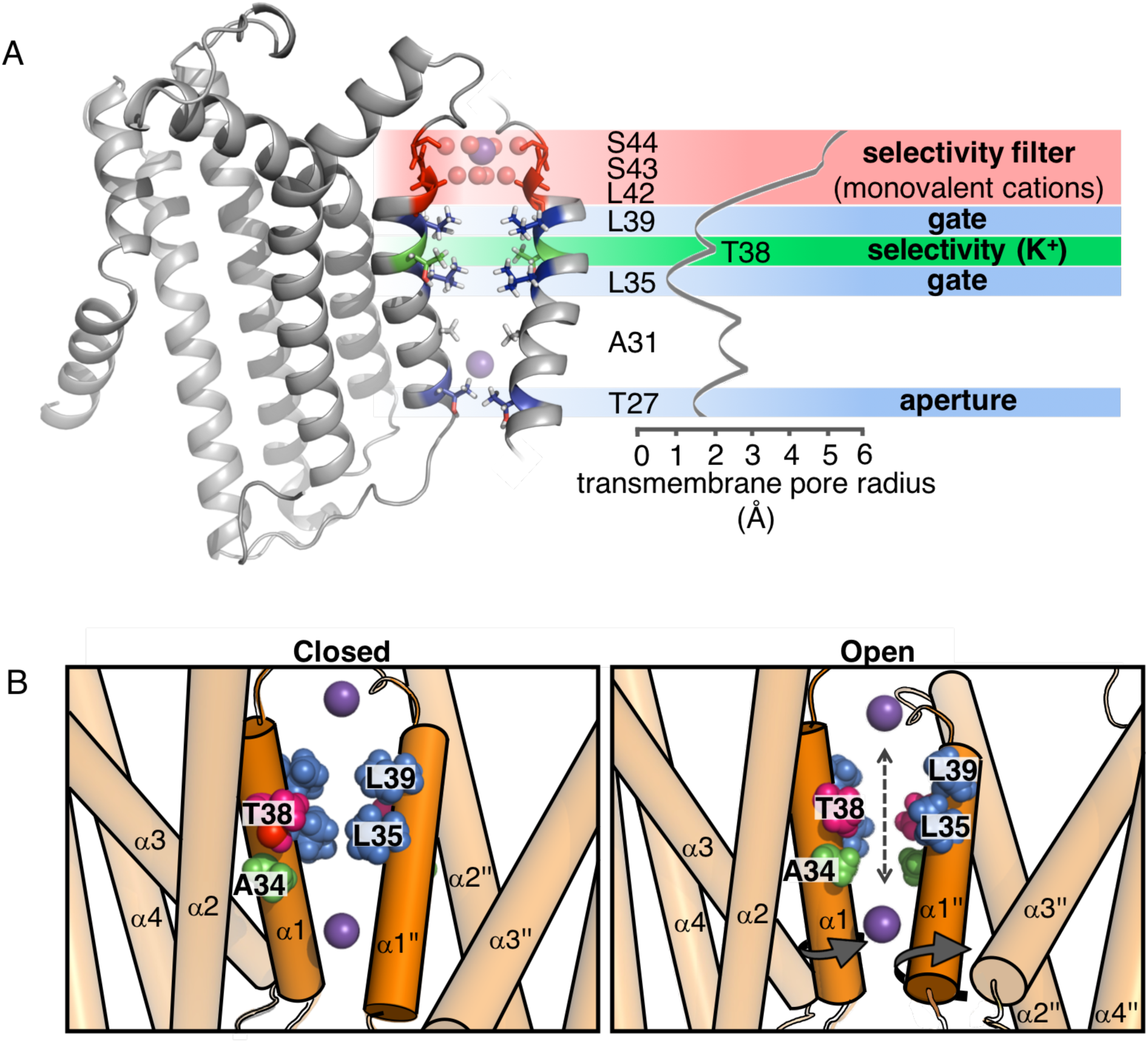
Functional layers and proposed mechanism for channel opening in TMEM175. **(A)** Functional layers in the MtTMEM175 pore. Two subunits are shown (right side only partly). Important residues and for comparison the HOLE analysis (see Fig. 3A) are indicated. Upper SF is shown in red, gate-residues in blue and inner SF in green. Thr^27^ is pore constricting in MtTMEM175, but is not conserved in other TMEM175 channels. **(B)** Schematic side view of MtTMEM175 in closed (left) and proposed conductive state (right). Key residues on helix 1 are shown as spheres. Two subunits are omitted for clarity. K^+^ ions (purple spheres) are positioned as in the structure. A clockwise rotation (viewed from intracellular) of 10-15° would widen the pore sufficiently for K^+^ ions to permeate (indicated by curved arrows in the right panel).

Threonine^38^ is also conserved in both repeats of vertebrate TMEM175 channels, but the K^+^ selectivity is about 10-20 times higher compared to the bacterial counterparts suggesting that additional features increase selectivity. Importantly, in vertebrate TMEM175 channels the two-fold symmetric pore may give rise to altered properties compared to the four-fold symmetric pore in the bacterial homologues. As proposed above, Ala^34^ in MtTMEM175 could face the pore upon rotation of helix 1. In repeat 1 but not 2 of all vertebrate TMEM175 channels, the corresponding residue is not an alanine but a serine, which could like Thr^38^ interact with permeating ions, to contribute additionally to selectivity (fig. S9A).

Our analysis of a bacterial TMEM175 channel has revealed ion binding sites and the architectural framework for ion selectivity in this protein family. On the one hand, TMEM175 channels are not as enigmatic as anticipated earlier but instead recapitulate classical elements of other ion channel families: Large hydrophobic residues acting as gates and polar contacts from side chains and the backbone to coordinate ions. On the other hand, it is remarkable that the selectivity is mediated by a cryptic threonine inside the pore that is only accessible in a conductive conformation, but not by the extracellular SF despite the obvious coordination of a K^+^ ion therein. Further, the conductive conformation must deviate substantially from the closed conformation, in order to be permeable to ions. This study provides insight into an alternative solution for conduction of K^+^ and expands our understanding on the biophysics of ion channels.

## Supporting information

## Acknowledgments

We are grateful to R. Dutzler (Univ. of Zürich) for supporting the project, providing genomic DNA, materials and lab infrastructure. We thank B. Blattmann and C. Stutz-Ducommun of the crystallization facility at Univ. of Zürich for support. S. Stefanic, P. Deplazes (Univ. of Zürich) are acknowledged for immunization of alpacas. We thank the staff team at the Swiss Light Source (PSI-Villigen) for excellent support. Calculations were performed at sciCORE (http://scicore.unibas.ch/) scientific computing center at University of Basel. We thank B. Dreier (Univ. of Zürich) and T. Sharpe (Univ. of Basel) for help with the MALLS experiment. At PSI/Villigen we thank T. Weinert, A. Prota and D. Ozerov for helpful discussions, and X. Li and R.A. Kammerer for support. We thank members of the Dutzler lab/Zürich and Maier lab/Basel for many fruitful discussions. A.M. and G.T. thank Henry Colecraft (Columbia University) for hospitality in his laboratory.

## Author contributions

J.D.B. and S.S. initiated and designed the project, cloned genes, purified and crystallized proteins, collected data at the synchrotron and analyzed data; S.S. designed the Nb-MBP fusion protein; Y.N. and J.D.B. generated the phage library; T.M. solved the initial crystal structure, R.P.J., T.M. and J.D.B. analyzed crystallographic data and carried out refinement; J.D.B: prepared final crystallographic models. T.S., A.M. and G.T. performed and analyzed electrophysiological experiments; S.S. and J.D.B. wrote the manuscript with input from all authors.

## Data and materials availability

Coordinates and structure factors have been deposited in the Protein Data Bank under accession codes XXXX, XXXX, XXXX, XXXX, XXXX, all data is available in the manuscript or supplementary materials.

