## Supplementary material for "Structural basis for ion selectivity in TMEM175 K^+^ channels"

**The Supplementary Materials includes:**

Materials and Methods
Supplementary Text
- -Design of the Nb-MBP fusion protein (macrobody) as a tool in structural biology
- -Comparison of the network of interactions connecting helices 1-3 and adjacent subunits in MtTMEM175 and CmTMEM175

Figs. S1 to S19
Table S1 to S2

### Materials and Methods

#### Cloning

Thirty TMEM175 genes were cloned from genomic DNA of various eubacteria. The genes were flanked by a 3C protease cleavage site, a myc-tag and a StrepTagII, either on the N- or C-terminus. The TMEM175 gene of *Marivirga tractuosa* (UniProt accession # E4TN31) was cloned from the strain DSM 4126. The TMEM175 cDNA of *Streptomyces collinus* (UniProt accession # S5VBU1) was synthesized by GenScript. For expression in MC1061 *E. coli*, TMEM175 genes were expressed from the FX-cloning plasmid pBXC3H (*50*) (Addgene # 47068) with a stop codon. For electrophysiology and expression in HEK293 cells, the TMEM175 genes were cloned without tags into the plasmids pcDXC3MS (*51, 52*) (Addgene #49030) followed by a stop-codon as well as into the vector pcDXC3GMS (*51, 52*) (Addgene #49031) (where EGFP was replaced by Venus-YFP using the *KpnI* sites) to obtain a C-terminally Venus-YFP tagged channel. Both, tagged and untagged versions yielded similar results. For stable cell lines, the expression cassette from pcDXC3MS containing the MtTMEM175 gene including a stop codon was shuttled to pcDNA5/TO (Thermo Fisher) vector using *HindIII* and *ApaI*. HEK293 Flp-In/T-Rex cells were transfected using FuGene (Promega) and selection was performed with 100 µg/ml hygromycin following the manufacturer's instructions. Expression was induced by addition of 3µg/ml tetracycline to the medium. For the selection of nanobodies, the MtTMEM175 gene was cloned into pBXC3H to purify the protein using a deca- histidine tag. An Avi-Tag for biotinylation was introduced by PCR at the C-terminus. Positive nanobodies were subcloned into the plasmid pBXNPHM3 (*52-54*) (Addgene #110099) for expression. C-terminally MBP (malE, *Escherichia coli K12*) tagged versions of nanobodies were generated by cloning nanobody genes and N-terminally truncated MBP genes into pBXNPHM3. The last four amino acids of MBP (RITK) were replaced with PGA. The resulting expression construct consists of a nanobody, a valine linker that connects the N-terminally truncated MBP, preceded by the pelB leader sequence, a deca- histidine tag, an MBP and a 3C protease cleavage site as depicted in fig. S3C. Mutant proteins were generated by site directed mutagenesis. All constructs were verified by Sanger sequencing.

#### Cell culture and transfection protocol

Membrane currents were recorded from cells, which were stably expressing MtTMEM175 or from cells transiently expressing the protein with similar results. For transient transfection low passage human embryonic kidney (HEK293T) cells were cultured in Dulbecco's modified Eagle's medium (Euroclone) supplemented with 10% fetal bovine serum (Euroclone), 100 IU/mL of penicillin, 100 µg/ml of streptomycin, and stored in a 37 °C humidified incubator with 5% $CO_2$. Transfections were performed with TurboFect Transfection Reagent (Thermo Scientific) according the producer protocol: The

MtTMEM175 gene inserted in pcDXC3MS was co-transfected with a plasmid containing green fluorescent protein (GFP) and incubated in dark.

#### Patch clamp recordings

One to two days after transfection, cells were dispersed by trypsin-EDTA treatment and seeded on 35-mm plastic petri dishes to allow single cell measurements. Green fluorescent cells were selected for patch clamp measurements. Membrane currents were recorded in whole cell configuration using an EPC9 or EPC10 patch-clamp amplifier (HEKA Electronics) controlled by the PatchMaster software (HEKA). Micropipettes with a resistance of about 2 MΩ were made from 1.5 mm thin-walled glass and fire-polished. The pipette solution contained (in mM) 150 KOH, 5 HCl, 10 HEPES, pH 7.4, titrated with methanesulfonic acid. The bath solution contained (in mM) 150 KOH, 1 $CaCl_2$, 1 $MgCl_2$, 10 TEA, 10 HEPES/KOH, pH 7.4, titrated with methanesulfonic acid. For measurements of selectivity $K^+$ was replaced by other cations of interest. Differences in osmolarity between pipette and bath solution were compensated by D-mannitol. In a standard voltage protocol the cell was clamped for 200 ms in 20 mV step protocol from holding voltage (0 mV, 100 ms) to test voltages between ±100 mV before returning to holding voltage (100 ms). The steady state current at the test voltages was measured during the final 20 ms of clamp steps.

#### Expression and purification of MtTMEM175

MC1061 *E.coli* cells harboring the C-terminally tagged MtTMEM175 gene were grown in terrific broth with 100 µg/ml ampicillin to an $OD_{600}$ of 0.5 at 37°C. Expression was induced with 0.02% Arabinose and continued over night at 18°C. Cells were harvested and resuspended in 150 mM NaCl, 50 mM Hepes-NaOH pH 7.6, 10% glycerol containing protease inhibitors (Complete, Roche), DNase I and 5 mM $MgCl_2$. Cells were lysed at 15000-25000 p.s.i. Cell debris was removed by centrifugation at 8000g for 30 minutes. Membranes in the supernatant were harvested by centrifugation using a 45 TI rotor (Beckmann) at 42000 r.p.m. for 1 h and resuspended in 250 mM KCl, 20 mM Hepes-NaOH pH 7.6, 15% glycerol. Extraction of the protein was carried out using 2% *n*-dodecyl-β-d-maltopyranoside (DDM, Anatrace) and protease inhibitors (Roche) for 1 h and subsequently centrifuged at 42000 r.p.m. using a 45 Ti rotor (Beckmann). The supernatant was incubated in batch with Strep-Tactin resin (Strep-Tactin Superflow high capacity, iba/Göttingen) for 1 h, washed with 150 mM KCl, 10 mM Hepes-NaOH pH 7.6, 10 % glycerol, 50 µg $ml^{-1}$ *E.coli* polar lipids (Avanti) and 0.03% DDM, and MtTMEM175 was eluted with the wash buffer containing 5 mM *d*-Desthiobiotin (Sigma-Aldrich). The protein was cleaved using HRV 3C protease and concentrated to 10-20 mg/ml using Amicon concentrators (Millipore) with a 100kDa cutoff. The MtTMEM175 protein was mixed with $Nb_{51H01-MBPs}$ in a molar ratio of 2.2– 2.5. For this, concentrated $Nb_{51H01-MBPs}$ was supplemented with 3mM maltose to keep MBP in the substrate-bound conformation. Subsequently DDM was added to 0.03 % and after that concentrated MtTMEM175 was added for complex formation. The mixture was left on ice for 30 minutes and applied to a Superdex 200 10/300 column (GE healthcare) equilibrated with 150 mM KCl, 5 mM Hepes-NaOH pH 7.6, 2.5 mM Maltose and 0.03 % DDM. The peak fractions were pooled and concentrated to 8-16 mg/ml for crystallization. All Steps were performed on ice or at 4 °C. Mutant proteins were purified in the same way.

#### Multi angle laser light scattering (MALLS) measurements

3C-protease cleaved MtTMEM175 protein was purified as described above except that the peak fraction after size exclusion chromatography was diluted to 35 μM (1mg/ml) before subjecting it to MALLS-SEC using a Superdex 200 10/300 column (GE healthcare) with an Agilent LC-1100 system coupled to an Optilab rEX refractometer (Wyatt Technology) and a miniDAWN 3-angle light-scattering detector (Wyatt Technology)). The SEC buffer contained 150 mM KCl, 10 mM Hepes-NaOH and 0.03 % DDM at pH 7.6 at RT. Data was analyzed with ASTRA software (Wyatt Technology).

#### Generation of nanobodies in Alpacas

Nanobodies against MtTMEM175 were raised in alpacas (*Vicugna pacos*) at the University of Zurich (now Nanobody Service Facility, NSF/UZH) as previously described (*52*). Briefly, alpacas were immunized four times with 14-day intervals by injecting 100 μg of purified MtTMEM175 protein at a concentration of 35 μM (in 150 mM KCl, 10 mM Hepes-NaOH pH 7.6, 0.03 % DDM, 15% glycerol) subcutaneously. A blood sample was used to generate lymphocyte cDNA by reverse transcription. Nanobody genes were cloned into a phagemid vector to create a phage library which was screened by biopanning against biotinylated MtTMEM175 immobilized on Neutravidin-coated plates. Biotinylation was performed as described using recombinant BirA enzyme (*53, 54*). Positive binders were identified using ELISA and subcloned into pBXNPHM3 for expression.

#### Nanobody purification

For expression of nanobodies in the vector pBXNPHM3, MC1061 *E. coli* cells were grown to an $OD_{600}$ of 0.75 at 37°C in terrific broth containing 100 μg/ml ampicillin. Protein expression was started by addition of 0.02% Arabinose and continued for 3.5 h at 37°C. Cells were harvested and resuspended in 150 mM NaCl, 50 mM Tris-HCl pH 8, 20 mM imidazole, 5 mM $MgCl_2$, 10% glycerol, 10 μg/ml DNAse I and protease inhibitors (Complete, Roche). Cells were lysed at 15000-25000 p.s.i. Cell debris was removed by centrifugation at 42000 r.p.m in a 45 Ti rotor. The supernatant was applied in batch to

NiNTA-resin for 1 h, washed with 150 mM KCl, 40 mM imidazole pH7.6, 10% glycerol and eluted with 150 mM KCl, 300 mM imidazole pH 7.6, 10% glycerol. The protein was cleaved over-night using HRV 3C protease during dialysis against 150 mM KCl, 10 mM Hepes-NaOH, 20 mM imidazole, pH 7.6, 10% glycerol. The MBP–$His_{10}$-fragment was removed by binding to NiNTA resin and the flow-through containing the nanobodies was concentrated (Amicon) and applied to a Superdex 200 column (GE healthcare) equilibrated in 150 mM KCl, 5 mM Hepes 7.6. The peak fractions were concentrated to 10-25 mg/ml before mixing with MtTMEM175 for complex formation. Nanobodies with a C-terminal MBP fusion were expressed and purified in the same way. Complex formation of purified nanobodies with MtTMEM175 was analyzed by SEC, where $Nb_{51H01}$ was identified as an MtTMEM175 binder with a 1:1 stoichiometry.

#### Crystallization of the MtTMEM175-$Nb_{51H01MBPs}$ complex

Expression and monodispersity of purified TMEM175 proteins in small scale was analyzed by SDS-PAGE and SEC. Several TMEM175 proteins were expressed at reasonable rates and eluted as monodisperse species from SEC. Expression was scaled up and we could crystallize several homologues readily. However, all of the crystallized proteins, including MtTMEM175, diffracted not beyond 20 Å, even after intense optimization of the crystallization conditions. To improve crystallization, we generated nanobodies against MtTMEM175 as described above. $Nb_{51H01}$, identified by ELISA and SEC, was used for complex formation with MtTMEM175 and this complex was subjected to crystallization. The best crystals of this complex diffracted not beyond 12-15Å. To improve crystallization further we decided to fuse MBP to the C-terminus of $Nb_{51H01}$ in order to increase possible crystal contacts and the chance for advantageous crystal lattices. We fused the Nb at the C-terminus with an N-terminally truncated MBP (starting at $Lys^6$ without the signal sequence) linked by a valine residue as depicted in figure S3. This resulted in the interfacial sequence PVTV**V***KLVIWIN* (Nb C-terminus underlined, linker in bold and MBP N-terminus in italics) and we named the construct $Nb_{51H01\text{-}MBPs}$ (see also Supplementary Text). A complex of $Nb_{51H01\text{-}MBPs}$ and MtTMEM175 was purified by SEC (fig. S1). Before subjecting the sample to SEC the mixture (MtTMEM175: $Nb_{51H01\text{-}MBPs}$ ratio of 1:2.2-2.5) was left on ice for 15-30 minutes and eluted in 150 mM KCl, 5 mM Hepes-NaOH, 2.5 mM maltose and 0.03% DDM. The fractions containing the complex were concentrated to 8-16 mg/ml and subjected to crystallization trials.

Prior to crystallization the purified MtTMEM175-$Nb_{51H01\text{-}MBPs}$ complex was mixed with *E.coli* polar lipids (Avanti) and with *n*-decyl-β-d-maltopyranoside (DM, Anatrace) to a final concentration of 100 µg/ml and 0.3% respectively. Best crystals were obtained in a condition composed of 100 mM Tris-

HCl pH 8.5, 150 mM NaCl, 150 mM $MgCl_2$ and 28- 30% PEG400 grown at 20°C. After 14 days they were dehydrated for 3-4 h using mother liquor with 36% PEG400, cryo-protected and flash-frozen in liquid propane or liquid nitrogen with similar results. The crystals giving the best datasets were additionally soaked in a cryo-protecting solution containing 5 mM $KPtCl_4$ followed by back-soaking in the cryo-protecting solution to get rid of excess platinum. For soaking in cesium and rubidium, 150 mM KCl in the cryo-protecting solution was replaced by 150 mM CsCl and 150 mM RbCl respectively. For the anomalous signal of zinc, crystals were soaked for 15 minutes in a cryo-protecting solution containing 0.5 mM $ZnSO_4$. The mutant MtTMEM175 with a T38A substitution was crystallized under similar conditions and crystals were flash frozen in liquid nitrogen.

##### Data collection and structure determination

X-ray diffraction data was collected on the X06SA beamline at the Swiss Light Source (SLS) of the Paul Scherrer Institute (PSI) equipped with an EIGER 16M detector (Dectris) at 100K. Data reduction was performed using XDS (*55*) and XSCALE (*56*). The resolution cut off was determined by $CC_{1/2}$ criterion (*57*). Crystals of MtTMEM175 in complex with $Nb_{51H01\text{-}MBPs}$ belong to space group $P42_12$ (a = 131.2 Å, b = 131.2 Å, c = 132.6 Å), with a solvent content of 64%. Best diffracting crystals were obtained after soaking in $KPtCl_4$, but no anomalous platinum signal was detected. For the native data set seven datasets from a single crystal were merged together. Phases were obtained by molecular replacement in PHASER (*58*), using the individual atomic coordinates of MBP (PDB ID: 1ANF) (*59*), and the nanobody Nb60 (PDB ID: 5JQH) (*60*). An initial round of model refinement was performed using REFMAC5 (CCP4 program suite) (*61*) (*62*), followed by density modification with Parrot (*63*) and automated model building by Buccaneer (*64*). The initial model was improved by iterative cycles of manual model building in Coot (*65*) and refined in Buster-TNT (*66*), yielding excellent geometry (Ramachandran favored/outliers: = 95.9%/0.0%) and $R_{work}/R_{free}$ values of 0.209/0.253 (Tables S1 and S2). Potassium ion positions were verified by the anomalous signal at high wavelengths ($\lambda$ = 2.02460 Å). Refinements using Buster-TNT indicated a high occupancy for $K^+$ at the position of $1K^+$ and lower occupancy for $K^+$ at $2K^+$. Thus, the presence of both, $K^+$ and $Na^+$, at $2K^+$ is possible. Native crystals were soaked with cesium, rubidium, and zinc and the respective ion position determined by the anomalous signal. The anomalous signal for cesium and rubidium ions was strong and identified their positions at the extracellular ion channel entrance (at $1K^+$). The anomalous signal for the data measured at the zinc K-edge ($\lambda$ = 1.24610 Å) was weak, suggesting only partial occupancy. The MtTMEM175 model and structure factors (code XXXX, XXXX, XXXX, XXXX, XXXX) have been deposited in the Protein Data Bank.

Regions not defined in the electron density include residues 1-3, 283-301 and 484-486 for the $Nb_{51H01-MBPs}$ expression construct, and residues 1-8 and 241-247 for MtTMEM175 (5.4 % in total). Residues 1-3 in the Nb correspond to the N-terminal remainder after 3C cleavage and would be GPS, and Residues 283-301 correspond to residues 166-184 in MBP (numbering without signal peptide) and residues 484-486 correlates to the end of MBP.

The program HOLE (*67*) was used to analyze the pore radius in the MtTMEM175 ion conduction pathway and the electrostatic potentials were calculated with the program APBS (*68*) with a grid spacing of 0.5 in a range of -5 to +5 kTe. Figure preparation was carried out in PyMOL (Schrödinger LLC). Maps were exported from Coot for use in PyMOL.

##### Projection of sequence conservation on the MtTMEM175 structure (fig. S19)

Fourteen bacterial TMEM175 sequences were aligned: Nine bacterial sequences obtained from a BLAST search using the sequence of hTMEM175, and four randomly chosen bacterial TMEM175 sequences were aligned with MtTMEM175. The conservation index from this multiple sequence alignment was calculated using AL2CO (*69*) and was then used to replace the values for the B-factors in the PDB file of MtTMEM175. Missing parts between the different sequences were assigned a value of -1 by default. More negative values as from the AL2CO conservation index output were set to -1. Conservation index was visualized in the MtTMEM175 structure using rainbow colors and with the minimum set to -1 (least conservation, blue) and the maximum set to 2.8 (maximal conservation, red). Sequences used for the alignment were: *Marivirga tractuosa, Lactobacillus rossiae, Mycobacterium sp., Humibacillus sp., Micromonospora chaiyaphumensis, Oscillatoria sp., Azospirillum brasilense, Niastella vici, Streptomyces collinus, Chryseobacterium sp., Streptacidiphilus carbonis, Fulvivirga imtechensis, Methylobacterium extorquens, Deinococcus geothermalis, Paenibacillus curdlanolyticus*.

#### Supplementary Text

##### Design of the Nb-MBP fusion protein (macrobody) as a tool in structural biology

Nanobodies were successfully used as crystallization chaperones for membrane proteins (*22, 53, 54*), similar to Fabs of monoclonal antibodies which have also led to high resolution structures in complex with membrane proteins. Despite many advantages over Fabs (*70*), Nbs lack behind in providing a large surface for crystal contacts and formation of crystal lattices like in the KcsA-Fab-complex or the EcClC-Fab-complex (*4, 71*) which greatly improved resolution compared to the uncomplexed proteins (*1, 72*)**.**

When crystallizing MtTMEM175 in complex with $Nb_{51H01}$ we could improve diffraction only slightly compared to uncomplexed MtTMEM175. This has persuaded us to design a chaperone that would be substantially different from the Nb itself in order to obtain a new set of crystallization conditions. For this we aimed to enlarge the Nb by a C-terminal fusion, opposite from the antigen binding site in order to increase the surface for crystallization. MBP has successfully been utilized as a fusion to the target protein for crystallization, mainly, if not exclusively, as an N-terminal moiety (*73*). We considered MBP as a fusion protein at the C-terminus of the Nb for several reasons: Its biochemical stability, its size of ~40 kDa (a protein of that size could already help to obtain phase information for many target proteins in crystallography and would become clearly visible in cryo-EM), its proven ability to aid crystallization, its natural occurrence in the periplasmic space (since Nbs are usually expressed in the periplasm) and the presence of preferred secondary structure elements for linkage to the Nb. The first secondary structure element of MBP is a beta strand starting at $Lys^6$ (numbering without signal peptide). Large parts of Nbs are also made up of beta strands. The course of the backbone in such a fusion protein that connects between the ends of two beta strands was more foreseeable compared to a fusion between a beta strand and an alpha helix. To minimize flexibility between the moieties we aimed to keep the distance between the beta strands of both moieties short by truncating the N-terminus of MBP. The first ten amino acids of the N-terminus of MBP after cleavage of the signal peptide are KIEEGKLVIW and in addition to full length MBP we made two truncated versions shortened by three and five amino acids at the N-terminus, respectively. As a universal linker between the two moieties we chose valine resulting in three different constructs named $Nb_{51H01\text{-}MBPl}$, $Nb_{51H01\text{-}MBPm}$ and $Nb_{51H01\text{-}MBPs}$ (long, medium and short) (fig. S3C). All could be expressed and the vast majority of tested Nb-MBP fusions (we tested ~20 different Nbs against numerous targets with the shortest design; data not shown) eluted as monodisperse peaks on SEC. The shortest construct using $Nb_{51H01}$ ($Nb_{51H01\text{-}MBPs}$) was the most promising design, since it was presumably the least flexible and a complex of this chaperone with MtTMEM175 resulted in a high-resolution structure described herein. We named this scaffold 'macrobody'. The structure revealed details about the linker region and the interface of the two moieties. Interestingly, the beta strand building the C-terminus of the Nb merges almost uninterrupted with the N-terminal beta strand of MBP. Only two residues, the linker-valine ($Val^{122}$ in the macrobody) and the starting amino-acid $Lys^6$ ($Lys^{123}$ in the macrobody) of the truncated MBP are not showing true beta-sheet architecture, but hydrogen bonds still involve the backbone of these two residues. Thus, numerous hydrogen-bonds, largely involving backbone amine hydrogens and carbonyl oxygens but also some interactions with side chains stabilize the connection between the Nb and MBP. These hydrogen bonds are also largely solvent inaccessible as they are shielded by mainly hydrophobic side chains, which increases the strength of the hydrogen bonds. (fig S3B). In addition, numerous bulky side chains at the N-terminal end of MBP and from loops at the C-terminal side of the Nb ensheath the linker, which likely reduces relative motions between the two moieties and contributes to the rigidity of the scaffold (fig S3E). This is also reflected by the relative temperature factors which are average in the linker region, similar to the Nb itself (fig S3F). The C-terminal side of Nbs is well conserved and known structures of Nbs align very well in the areas that come close to the MBP moiety, including also the position of the very C-terminus. This explains why this scaffold could be used with a number of different specific Nbs and is therefore of high general applicability. With this macrobody we have thus generated a very useful scaffold with excellent geometry, good biochemical properties and uncomplicated molecular biology that can be easily adapted to other Nbs. This modification of Nbs expands the potential of these binders in crystallization. The emergence of synthetic Nb-libraries and display techniques reduces the time for generation of Nbs greatly and allows to circumvent animal immunization, making Nbs also accessible at lower budgets (*74, 75*). Nbs have become wide spread tools in membrane protein biochemistry, and macrobodies could improve the success rate in crystallization, also for targets that resisted so far to crystallize despite the utilization of Nb chaperones. Furthermore, due to the large mass contribution of macrobodies, molecular replacement can be more readily used to solve a structure, as done in this study. Another technique where macrobodies could be very promising tools, is electron cryo-microscopy. Future hard- and software developments in cryo-EM will certainly allow for structural analysis of ever smaller proteins (*76-79*). However, enlarging binders will still be advantageous for structure determination of difficult targets, such as small membrane proteins lacking extramembrane domains. Here, Nbs are generally too small to be of much help in single particle classification. Fabs have however been utilized successfully to enlarge membrane proteins for this purpose (*80-82*) but are more complex to obtain, have a lower propensity to be conformation specific and are difficult to engineer. The Nb-MBP fusion protein or macrobody might show some flexibility in solution (which still needs to be tested) and flexibility would have to be largely abolished for cryo-EM applications. Fabs exhibit some intrinsical flexibility, and efforts have been attempted to tackle this problem (*83*). Clearly, the ease of engineering and expression of Nb-MBP fusion proteins makes this undertaking less cumbersome. Albeit a single connecting peptide chain would intuitively imply flexibility in the Nb-MBP fusion protein, the above-mentioned features of the structure suggest a relatively rigid molecule. Macrobodies hold great promise to be competitive to Fabs, which is already given with respect to its size (fig. S3). Further, the high content of alpha-helical secondary structure elements in MBP is increasing contrast in cryo-EM micrographs as compared to beta-sheet rich Fabs. Recent approaches to increase the size of small proteins for cryo-EM underline the demand for enlarging binders in this field of structural biology (*84*), and macrobodies could thus become a very useful addition to the already rich repertoire of Nb-applications.

#### Comparison of the network of interactions connecting helices 1-3 and adjacent subunits in MtTMEM175 and CmTMEM175

TMEM175 channels feature only very few highly conserved residues in the primary sequence (fig. S9B). When viewed from the intracellular side it becomes apparent that most of these conserved residues are essential for positioning of helices 1-3 towards each other and linking adjacent subunits in bacterial and vertebrate TMEM175 channels (figs. S13 and S14) by forming a network of mainly polar interactions. In MtTMEM175, $Arg^{24}$ is interacting with $His^{77}$ (from the adjacent subunit) perpendicular to the plane of the imidazole side chain. The highly conserved $Asp^{30}$ (part of the FSD motif and from the same adjacent subunit as the interacting $His^{77}$), is also interacting with $Arg^{24}$. $Arg^{24}$ is thus connecting to helix 1 and 2 of the neighboring subunit. Mutating $Arg^{24}$ to alanine or lysine resulted in increased amount of non-tetrameric protein in SEC, supporting a role of this residue in tetramer assembly (fig. S15), which makes it difficult to study additional functions of this residue. In the CmTMEM175 model, this particular arginine ($Arg^{12}$) is not bound to the corresponding histidine ($His^{73}$), but to the following residue ($His^{74}$) (figs. S13 and S14B) via a cation-π interaction (*27*). The position of the second histidine ($His^{74}$) is relatively conserved, and can be found also in hTMEM175 in the second repeat (fig. S9B). In MtTMEM175 it is a tyrosine in difference to most TMEM175 channels. Cation-π-stacks of arginine and histidine occur in protein structures but the interaction is pH-dependent (*85*). Positively charged histidines generally repel arginine side chains. When looking at the $2F_o$-$F_c$ map we could not see clear electron density in the CmTMEM175 structure for this arginine and even negative difference density in the $F_o$-$F_c$ map (fig. S14B). We thus conclude that $Arg^{12}$ is unbound. Importantly, CmTMEM175 was crystallized at pH 4.6 as opposed to pH 8.5 in MtTMEM175, and thus these solvent-accessible histidines are probably protonated, causing repulsion of the arginine. The hydrogen bond network between $His^{77}$, $Asn^{95}$ and $Ser^{29}$ is present in both structures (fig. S13). The π-stack between $Phe^{16}$ and $Trp^{70}$ of the adjacent subunit of CmTMEM175 is weaker in MtTMEM175 (here the two sidechains of $Phe^{28}$ and $Trp^{74}$ are angled towards each other). Instead $Trp^{74}$ is interacting with $Asp^{30}$ in MtTMEM175 whereas the larger distances of the corresponding pair $Asp^{18}$ and $Trp^{70}$ in CmTMEM175 indicate a weaker interaction. Whether these differences as well as the before mentioned $3_{10}$-helix are linked to the respective interactions of $Arg^{12}$ in CmTMEM175 and $Arg^{24}$ in MtTMEM175 is currently not clear. In CmTMEM175, the N-terminal end of helix 1 is tilted slightly away from the adjacent subunit, thus widening the intracellular pore entrance. Also, helix 2 and 3 in CmTMEM175 tilt outwards in comparison to MtTMEM175. As a consequence, the CmTMEM175 structure appears more open at the intracellular side, but nevertheless is representing also a non-conductive conformation (*27*). If this conserved arginine plays a role in opening and closing of the channel and if this is linked to the intracellular pH will have to be addressed in future experiments. A link between the intracellular pH and the open probability would have profound consequences for the understanding of the functional role of TMEM175 channels, both in bacteria and animals.

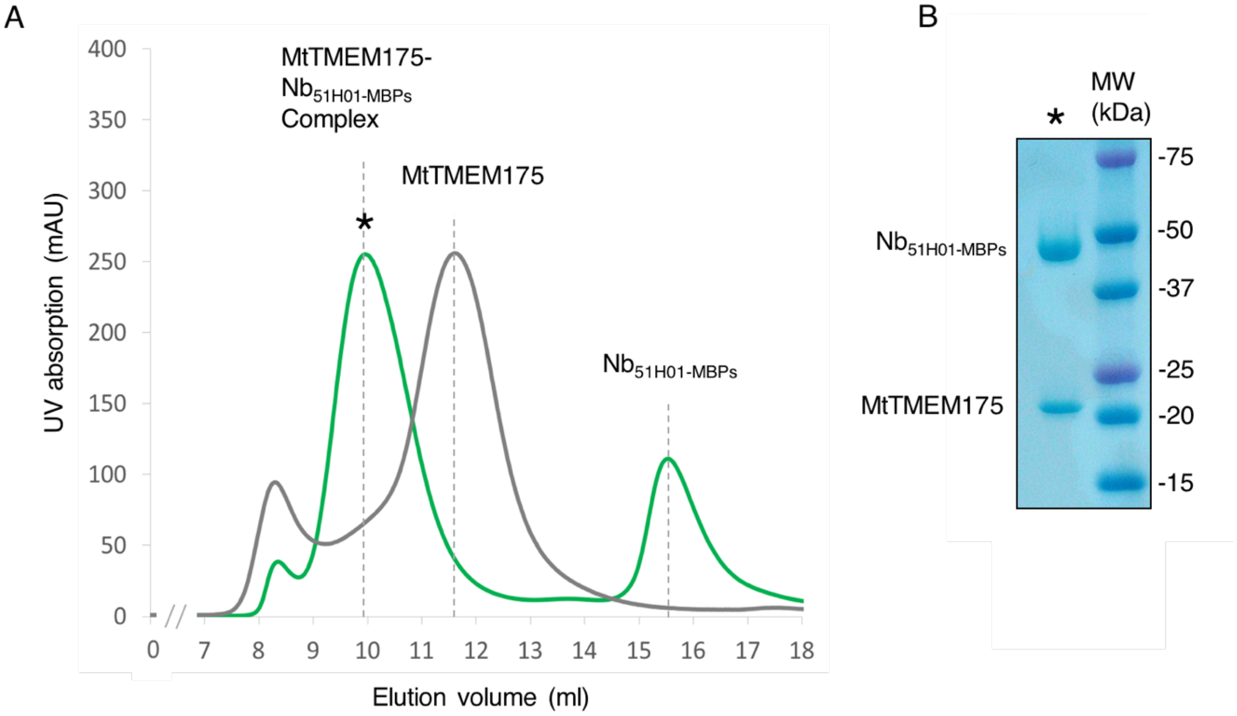


**Fig. S1. MtTMEM175 Nb$_{51H01\text{-}MBPs}$ complex formation. (A)** Size exclusion chromatogram from a Superdex 200 10/300 column showing MtTMEM175 (grey) and MtTMEM175-Nb$_{51H01\text{-}MBPs}$ complex (green). Excess Nb$_{51H01\text{-}MBPs}$ is eluting at larger elution volume. The peak height of MtTMEM175 is normalized to the peak height of the complex. **(B)** Coomassie-stained SDS-PAGE gel of the peak fraction (marked with an asterisk in **(A)**) indicating complex formation of MtTMEM175 with Nb$_{51H01\text{-}MBPs}$. The molecular weight marker (MW) is shown on the right.

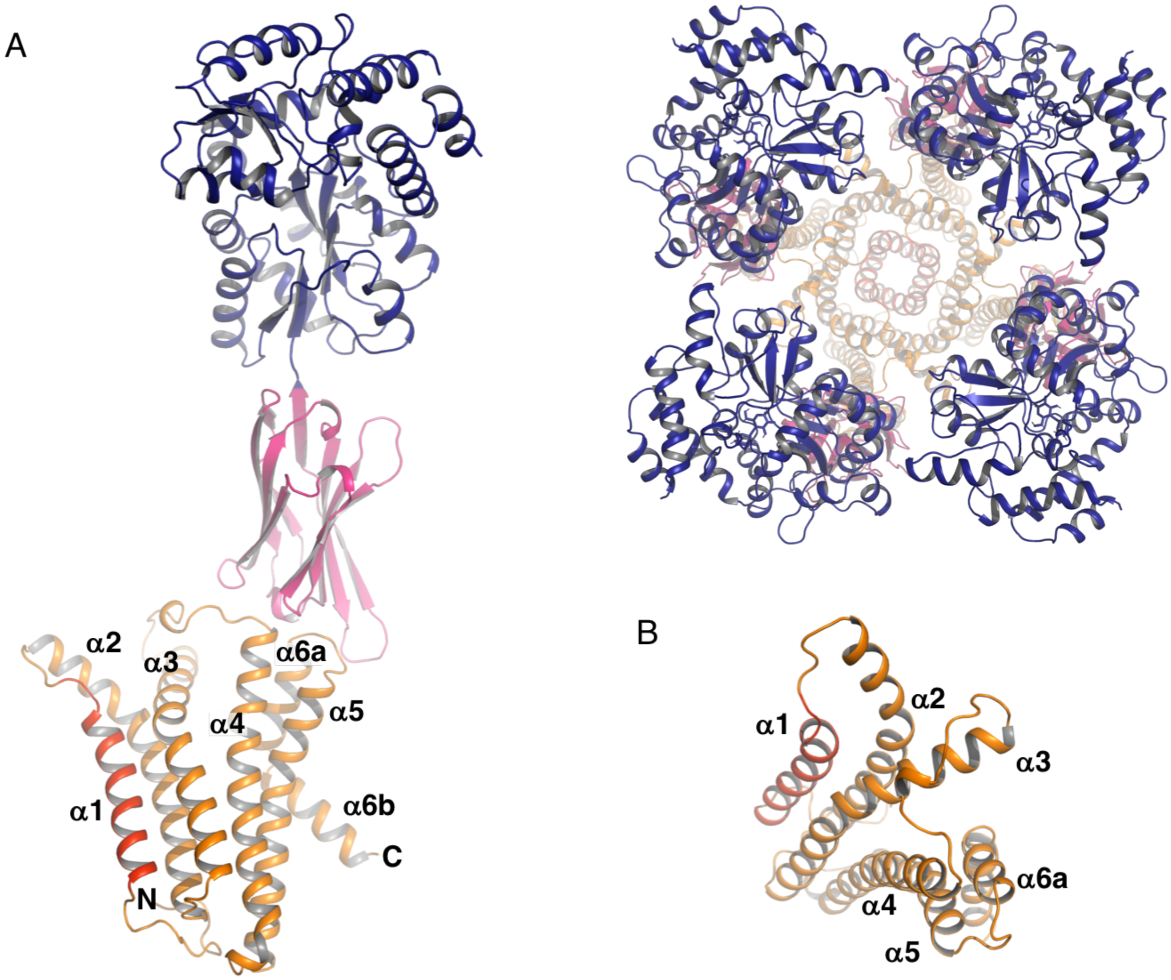


**Fig. S2. Structure of the complex. (A)** Side view of a single subunit (left) and top view (right) of a tetramer of the MtTMEM175- $Nb_{51H01\text{-}MBPs}$ complex in cartoon representation. **(B)** Top view of a single subunit of MtTMEM175. The Nb and the MBP moiety are colored in pink and dark blue, respectively, while MtTMEM175 is displayed in orange with the pore-lining helix (α1) in red. The numbering of transmembrane helices in MtTMEM175 is indicated.

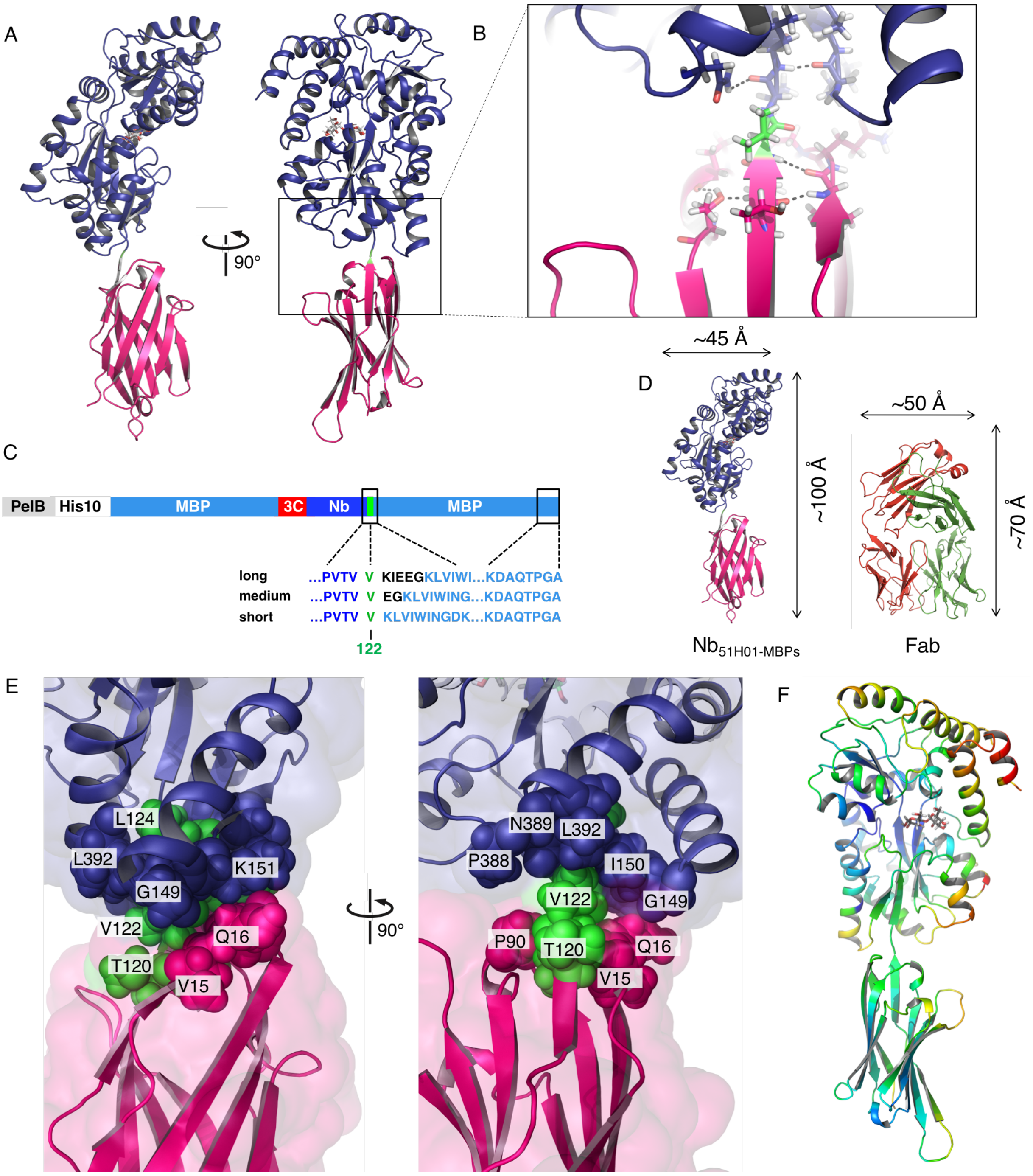


**Fig. S3**. **Structure of the $Nb_{51H01\text{-}MBPs}$ fusion protein (macrobody). (A)** Different side views. The bound maltose is shown as sticks in grey. **(B)** Close-up view of the linker region between the Nb and MBP moiety with hydrogen bonds indicated as black dashed lines. Valine$^{122}$ introduced as linker between the Nb and MBP is shown in green. **(C)** Expression construct. Valine$^{122}$ between the Nb and MBP moieties is highlighted in green. The short construct was used in this study and represents the scaffold of macrobodies. PelB: PelB leader sequence, His10: deca histidine-tag, MBP: maltose binding protein, 3C: 3C protease cleavage site, Nb: nanobody. **(D)** Dimensions of the $Nb_{51H01\text{-}MBPs}$ fusion protein and a Fab (anti-KcsA, PDB entry 5J9P) in comparison. The Nb and the MBP are colored pink and dark blue, respectively. **(E)** Close-up views of the linker region (orientation as in **(A)**). The connecting strand between MBP and Nb is colored in green (sequence $TVV^{122}KL$) and displayed as spheres, and parts belonging to MBP and Nb are shown in blue and pink, respectively. Residues ensheathing the linker are shown in sphere representation and labelled (numbering refers to the fusion protein). **(F)**. Relative B-factors in the macrobody in spectrum colors. Blue: lowest B-factors, red: highest B-factors.

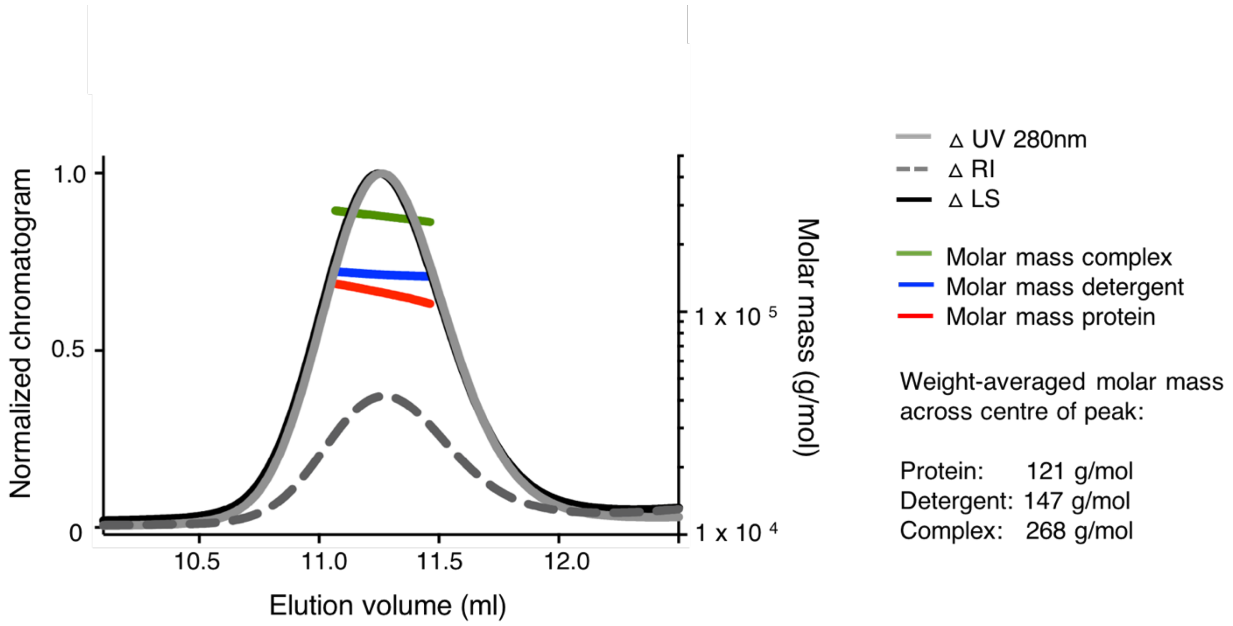


**Fig. S4. Characterization of the oligomeric state of MtTMEM175.** Multi angle laser light scattering (MALLS) coupled to a Superdex 200 (10/300) column. MtTMEM175, after cleavage with 3C protease, was prepared at a concentration of 35µM (1 mg/ml) and 50 µg were injected. The molar mass is indicated: red, MtTMEM175, blue, detergent micelle of DDM, green, MtTMEM175 with DDM detergent micelle. The calculated molecular weights of one subunit of MtTMEM175 and the tetramer are 29 kDa and 116 kDa respectively. RI: refractive index, LS: light scattering.

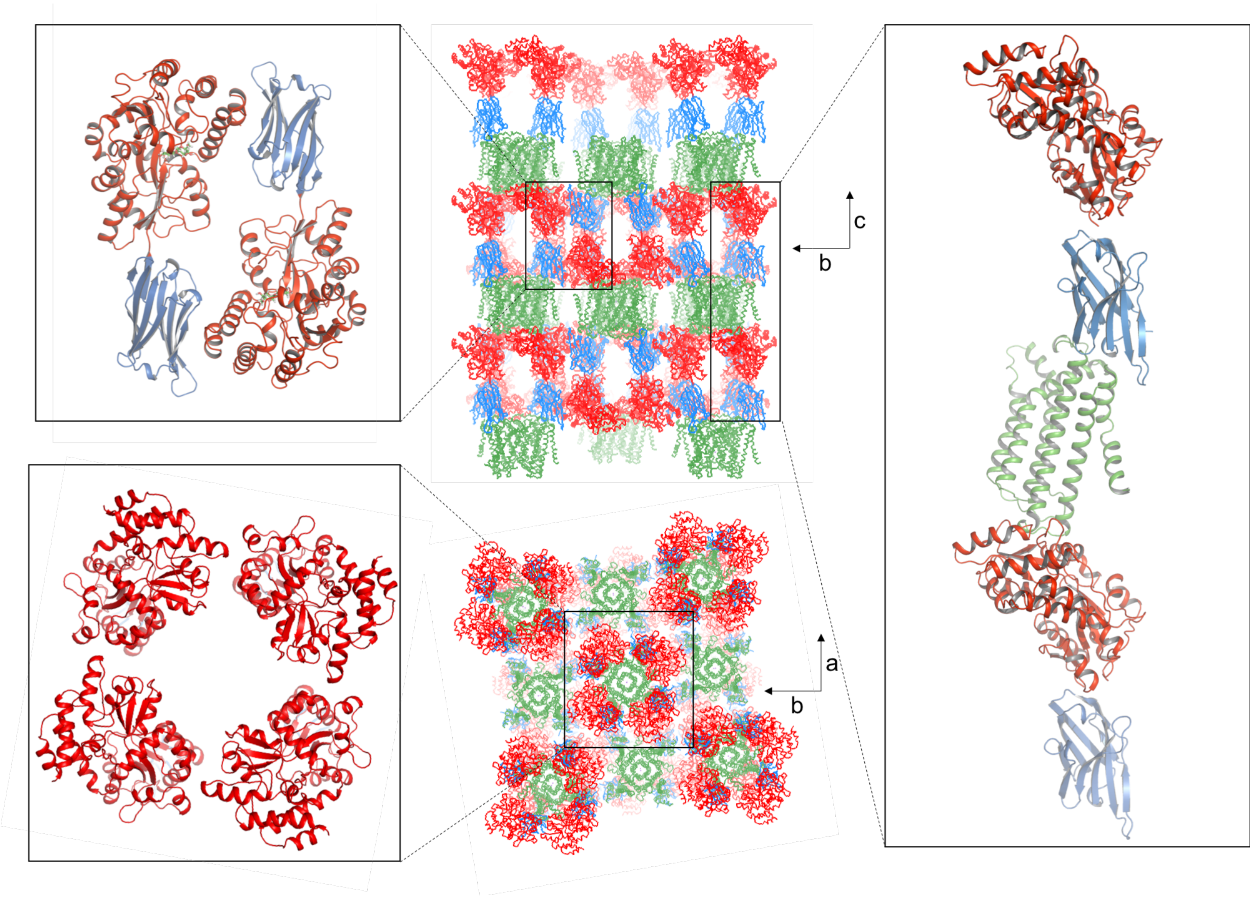


**Fig. S5. Crystal lattice in the crystal of the MtTMEM175-Nb$_{51H01\text{-}MBPs}$ complex.** Crystal contacts with the MtTMEM175 tetramer are only mediated by Nb$_{51H01\text{-}MBPs}$ (right box). Additional crystal contacts can be seen between two antiparallel Nb$_{51H01\text{-}MBPs}$ molecules (left upper box) and between MBP moieties (left lower box). There are no crystal contacts between individual MtTMEM175 tetramers. MtTMEM175 is shown in green, the Nb in blue and MBP in red. View along a and c axes.

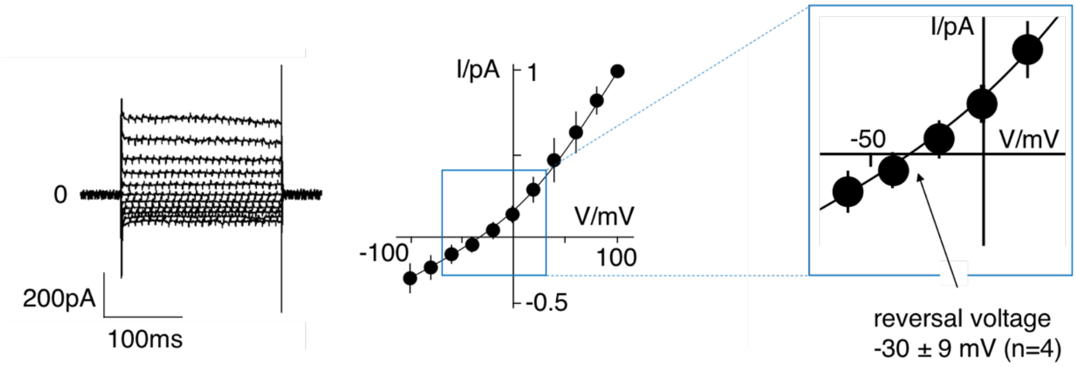


**Fig. S6. $K^+$ selectivity of ScTMEM175.** Current responses of HEK293 cells expressing ScTMEM175 to voltage steps between ±100 mV (left). Cells were measured with 150 mM $Na^+$ and 150 mM $K^+$ in the bath and pipette solution, respectively. Mean steady state I/V relation (±s.d.) from 4 cells measured under the same condition. Data were normalized to currents at +100 mV. The mean reversal voltage of the 4 cells is -30±9 mV.

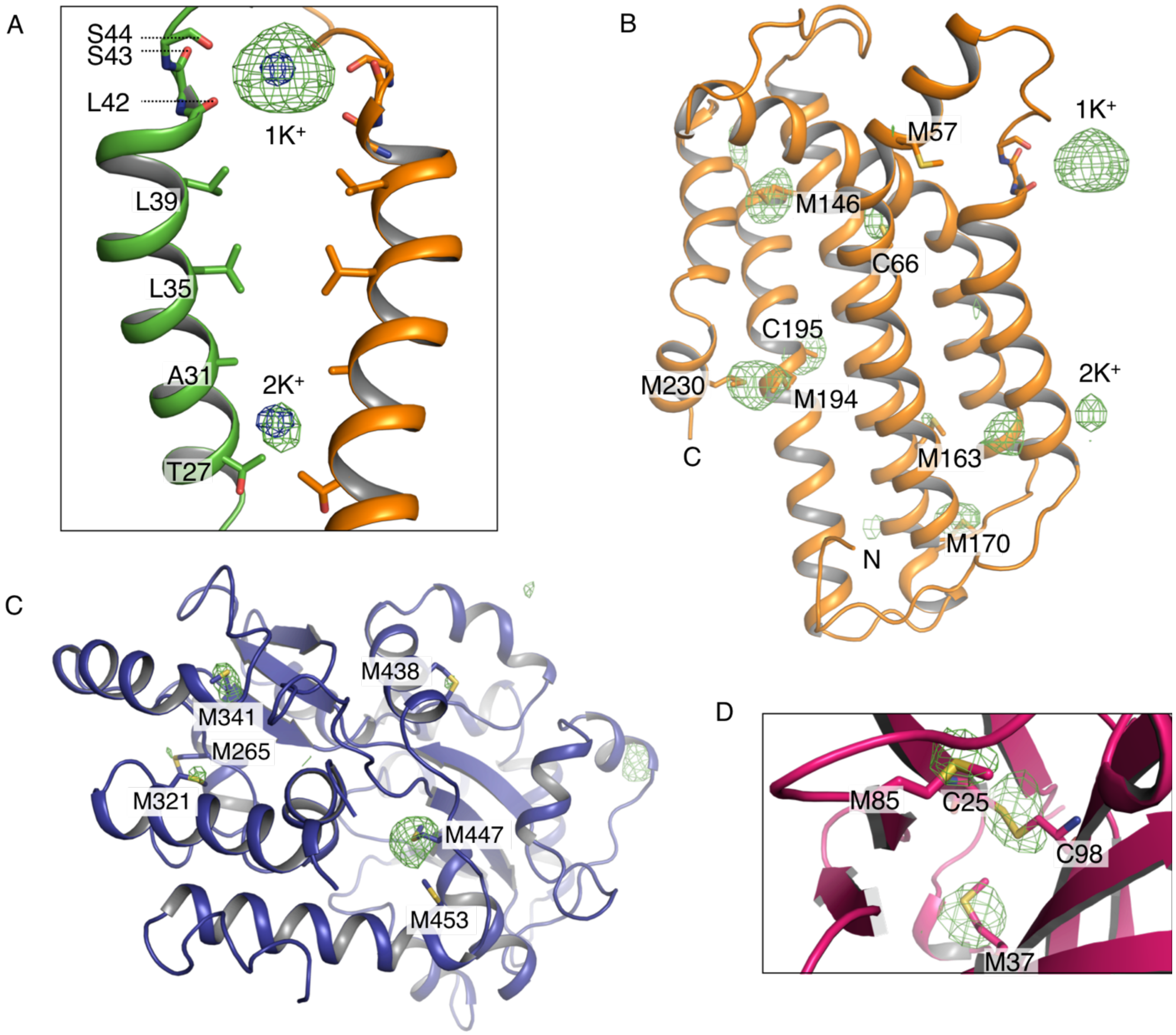


**Fig. S7. Anomalous difference electron density in the MtTMEM175-Nb$_{51H01-MBPs}$ complex. (A)** Verification of $K^+$ ions at positions $1K^+$ at the extracellular entrance and $2K^+$ within the pore close to the intracellular entrance. The signal was detected at a wavelength of 2.02460 Å (Table S1, K/S). The anomalous difference density map at +3 σ is shown as green mesh (at 3.5 Å, blurred with b=165). For comparison, the $2F_o$-$F_c$ electron density from the best dataset (at 2.4 Å, contoured at 1.8 σ, sharpened with b=-25) is illustrated as blue mesh at the respective positions of $1K^+$ and $2K^+$ (see also Table S1, native). A side view of helix 1 of two opposing subunits is depicted. **(B)** View on MtTMEM175 (only one subunit is shown), **(C)** MBP and **(D)** Nb with the anomalous difference density superimposed on the model with same settings as in **(A)**. In **(D)** a close-up view on the Nb is shown. All Cys and Met residues of the complex are labeled and overlaid well with the anomalous signal detected at 2.02460 Å, confirming the correct register of the structure.

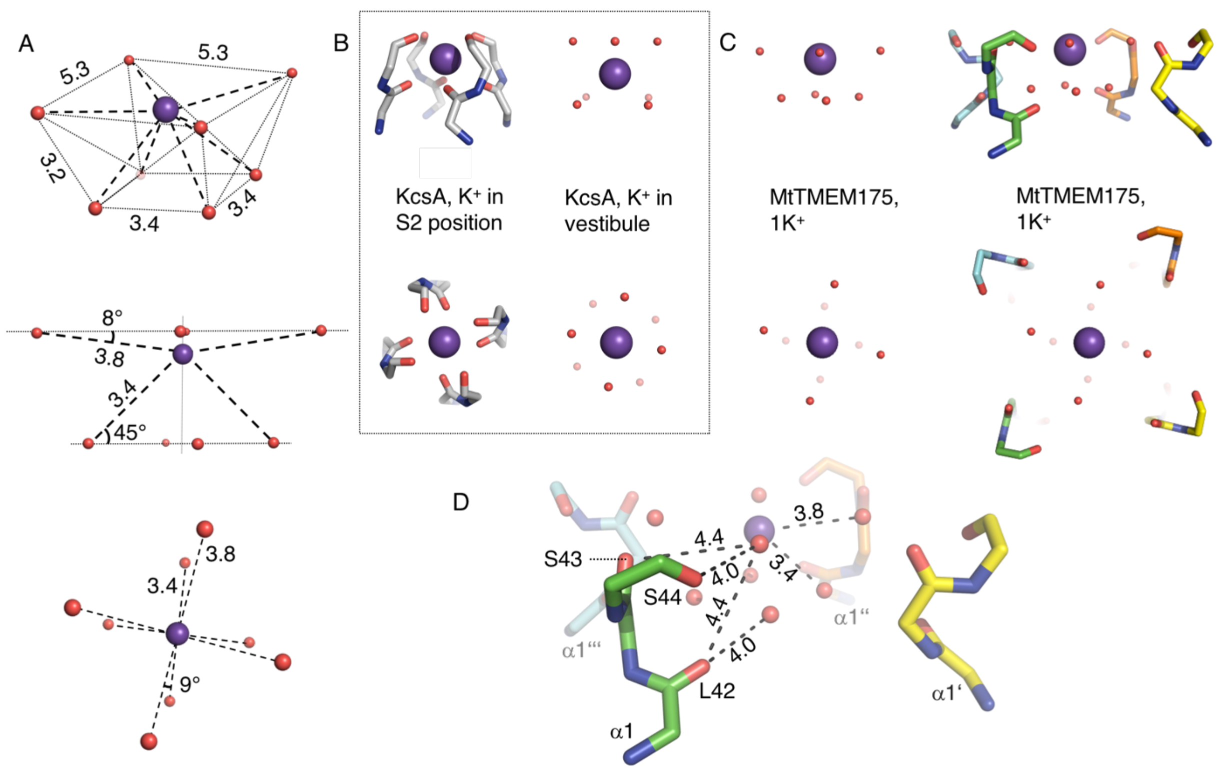


**Fig. S8. Coordination of $K^+$ in MtTMEM175 and KcsA. (A)** Geometry and dimensions of the hydrated $K^+$ in MtTMEM175. Angles and atom-to-atom distances are indicated in degrees (°) and Å, respectively. Views from the side and from extracellular. **(B)** Coordination of $K^+$ in KcsA by backbone oxygens (left, PDB entry 1K4C, S2 position) and geometry of a hydrated $K^+$ ion in the KcsA vestibule in proximity to the SF (right, PDB entry 1K4C). **(C)** Geometry of the hydrated $K^+$ ion at $1K^+$ and coordination of this hydrated $K^+$ by backbone oxygens of residues $Leu^{42}$, $Ser^{43}$ and $Ser^{44}$ in MtTMEM175. **(D)** Interactions of the backbone oxygens with the hydrated $K^+$ ion. Atom-to-atom distances are indicated in Å. The water molecules and $K^+$ ions are shown as spheres with reduced size for clarity.

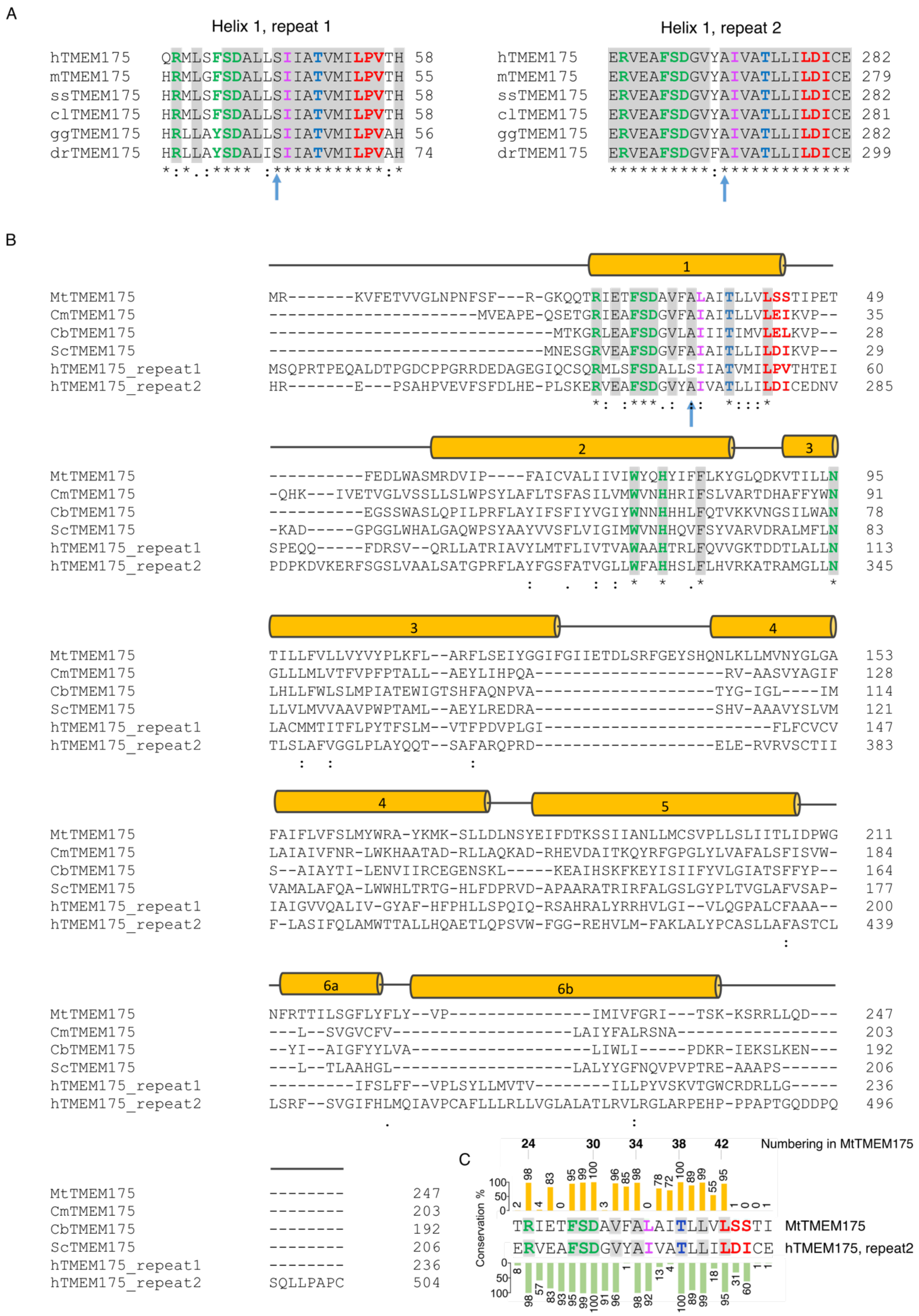
A
Helix 1, repeat 1
hTMEM175 QRMLSFSDALLSIIATVMILPVTH 58
mTMEM175 HRMLGFSDALLSIIATVMILPVTH 55
ssTMEM175 HRMLSFSDALLSIIATVMILPVTH 58
clTMEM175 HRMLSFSDALLSIIATVMILPVTH 58
ggTMEM175 HRLLAYSDALLSIIATVMILPVAH 56
drTMEM175 HRLLAYSDALISIIATVMILPVAH 74
Helix 1, repeat 2
hTMEM175 ERVEAFSDGVYAIVATLLILDICE 282
mTMEM175 ERVEAFSDGVYAIVATLLILDICE 279
ssTMEM175 ERVEAFSDGVYAIVATLLILDICE 282
clTMEM175 ERVEAFSDGVYAIVATLLILDICE 281
ggTMEM175 ERVEAFSDGVYAIVATLLILDICE 282
drTMEM175 ERVEAFSDGVFAIVATLLILDICE 299
B
MtTMEM175 MR-------KVFETVVGLNPNFSF---R-GKQQTRIETFSDAVFALAITLLVLSSTIPET 49
CmTMEM175 ----------------------MVEAPE-QSETGRIEAFSDGVFAIAITLLVLEIKVP-- 35
CbTMEM175 ------------------------------MTKGRLEAFSDGVLAIIITIMVLELKVP-- 28
ScTMEM175 -----------------------------MNESGRVEAFSDGVFAIAITLLILDIKVP-- 29
hTMEM175_repeat1 MSQPRTPEQALDTPGDCPPGRRDEDAGEGIQCSQRMLSFSDALLSIIATVMILPVTHTEI 60
hTMEM175_repeat2 HR-------E---PSAHPVEVFSFDLHE-PLSKERVEAFSDGVYAIVATLLILDICEDNV 285
MtTMEM175 ----------FEDLWASMRDVIP----FAICVALIIVIWYQHYIFFLKYGLQDKVTILLN 95
CmTMEM175 -QHK---IVETVGLVSSLLSLWPSYLAFLTSFASILVMWVNHHRIFSLVARTDHAFFYWN 91
CbTMEM175 ----------EGSSWASLQPILPRFLAYIFSFIYVGIYWNNHHHLFQTVKKVNGSILWAN 78
ScTMEM175 -KAD-----GPGGLWHALGAQWPSYAAYVVSFLVIGIMWVNHHQVFSYVARVDRALMFLN 83
hTMEM175_repeat1 SPEQQ-----FDRSV--QRLLATRIAVYLMTFLIVTVAWAAHTRLFQVVGKTDDTLALLN 113
hTMEM175_repeat2 PDPKDVKERFSGSLVAALSATGPRFLAYFGSFATVGLLWFAHHSLFLHVRKATRAMGLLN 345
MtTMEM175 TILLFVLLVYVYPLKFL--ARFLSEIYGGIFGIIETDLSRFGEYSHQNLKLLMVNYGLGA 153
CmTMEM175 GLLLMLVTFVPFPTALL--AEYLIHPQA--------------------RV-AASVYAGIF 128
CbTMEM175 LHLLFWLSLMPIATEWIGTSHFAQNPVA-------------------TYG-IGL----IM 114
ScTMEM175 LLVLMVVAAVPWPTAML--AEYLREDRA-------------------SHV-AAAVYSLVM 121
hTMEM175_repeat1 LACMMTITFLPYTFSLM--VTFPDVPLGI------------------------FLFCVCV 147
hTMEM175_repeat2 TLSLAFVGGLPLAYQQT--SAFARQPRD-------------------ELE-RVRVSCTII 383
MtTMEM175 FAIFLVFSLMYWRA-YKMK-SLLDLNSYEIFDTKSSIIANLLMCSVPLLSLIITLIDPWG 211
CmTMEM175 LAIAIVFNR-LWKHAATAD-RLLAQKAD-RHEVDAITKQYRFGPGLYLVAFALSFISVW- 184
CbTMEM175 S--AIAYTI-LENVIIRCEGENSKL-----KEAIHSKFKEYISIIFYVLGIATSFFYP-- 164
ScTMEM175 VAMALAFQA-LWWHLTRTG-HLFDPRVD-APAARATRIRFALGSLGYPLTVGLAFVSAP- 177
hTMEM175_repeat1 IAIGVVQALIV-GYAF-HFPHLLSPQIQ-RSAHRALYRRHVLGI--VLQGPALCFAAA-- 200
hTMEM175_repeat2 F-LASIFQLAMWTTALLHQAETLQPSVW-FGG-REHVLM-FAKLALYPCASLLAFASTCL 439
MtTMEM175 NFRTTILSGFLYFLY--VP---------------IMIVFGRI---TSK-KSRRLLQD--- 247
CmTMEM175 ---L--SVGVCFV-------------------LAIYFALRSNA----------------- 203
CbTMEM175 --YI--AIGFYYLVA-------------------LIWLI-----PDKR-IEKSLKEN--- 192
ScTMEM175 ---L--TLAAHGL-------------------LALYYGFNQVPVPTRE-AAAPS------ 206
hTMEM175_repeat1 ---------IFSLFF--VPLSYLLMVTV--------ILLPYVSKVTGWCRDRLLG----- 236
hTMEM175_repeat2 LSRF--SVGIFHLMQIAVPCAFLLLRLLVGLALATLRVLRGLARPEHP-PPAPTGQDDPQ 496
MtTMEM175 -------- 247
CmTMEM175 -------- 203
CbTMEM175 -------- 192
ScTMEM175 -------- 206
hTMEM175_repeat1 -------- 236
hTMEM175_repeat2 SQLLPAPC 504
C
24 30 34 38 42 Numbering in MtTMEM175
Conservation %
TRIETFSDAVFALAITLLVLSSTI MtTMEM175
ERVEAFSDGVYAIVATLLILDICE hTMEM175, repeat2

**Fig. S9. Sequence conservation in TMEM175 proteins. (A)** Sequence alignment of helix 1 in vertebrate TMEM175 proteins. h: human, m: mouse, ss: *Sus scrofa*, cl: *Canis lupus*, gg: *Gallus gallus*, dr: *Danio rerio*. **(B)** Comparison of sequences from characterized TMEM175 family members. Mt: *Marivirga tractuosa*, Cm: *Chamaesiphon minutus,* Cb: *Chryseobacterium sp.*, Sc: *Streptomyces collinus,* and h: human. For hTMEM175 both repeats were included in the alignment. Transmembrane helices from the MtTMEM175 structure are indicated as cylinders with corresponding numbering (see also fig. S2). In **(A)** and **(B)**: Identical amino acids are marked in grey. Residues forming the cation SF at the extracellular end of the pore are highlighted in red. Residues participating in a conserved hydrogen bond network between helices 1-3 are colored in green. The selectivity filter deeper in the pore ($Thr^{38}$) is marked in blue. The constriction point ($Leu^{35}$) is indicated in magenta. Blue arrows mark the position of a highly conserved serine and alanine in repeat 1 and 2 of vertebrate TMEM175 proteins, respectively. **(C)** A BLAST search within the bacterial phyla was done with the sequence of hTMEM175 and the first 100 hits were aligned against each other. The conservation between helix 1 of the bacterial homologs with helix 1 of MtTMEM175 (top) and hTMEM175 (repeat 2, bottom) is given in percent in a bar chart. The color code is the same as in **(A)** and **(B)** except that residues of MtTMEM175 and hTMEM175 exhibiting a conservation of ≥ 95% with the bacterial sequences are marked in grey in the two sequences, highlighting the most conserved residues within helix 1 in the TMEM175 family.

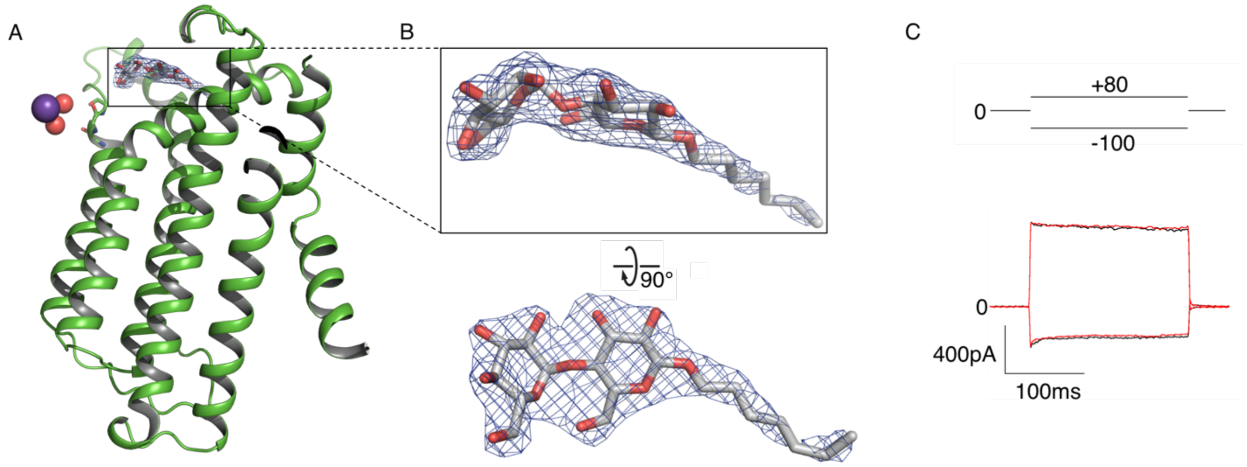


**Fig. S10. Electron density of a putative maltoside. (A)** Residual density on top of the channel was attributed to a detergent molecule (DDM or DM). The $2F_O$-$F_C$ density (at 2.4 Å, contoured at 1.5 σ after sharpening with b=-40, blue) is displayed. Only one subunit is shown. **(B)** Close-up views on the maltoside with overlaid $2F_O$-$F_C$ density. **(C)** Current responses of HEK293 cells expressing MtTMEM175 to voltage steps from 0 mV to +80/-100 mV before (black) and after (red) adding 10 mM maltose to the bath solution with 150 mM $Na^+$.

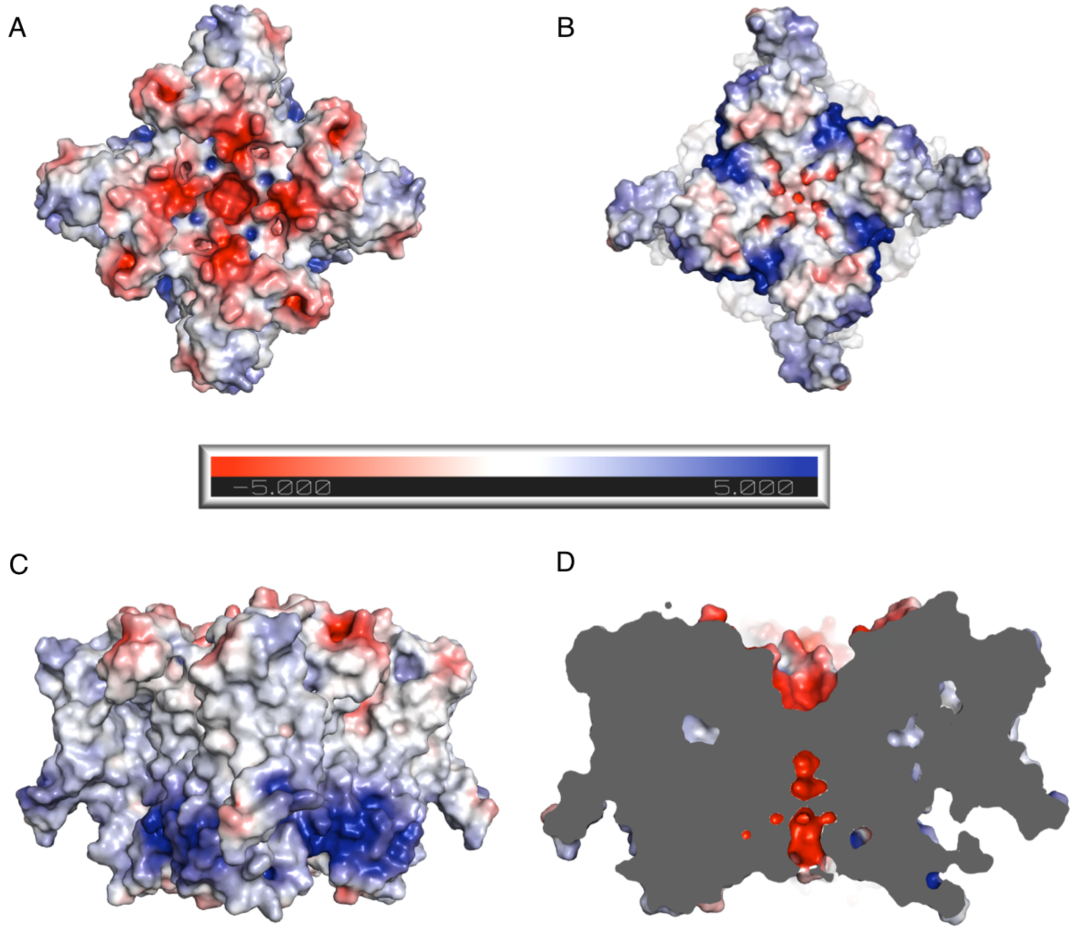


**Fig. S11. Illustration of the surface electrostatic potential in MtTMEM175.** View on the protein surface from extracellular **(A)**, intracellular **(B)** and from the side **(C)** and **(D)**. In **(D)** a cross section through the pore is shown.

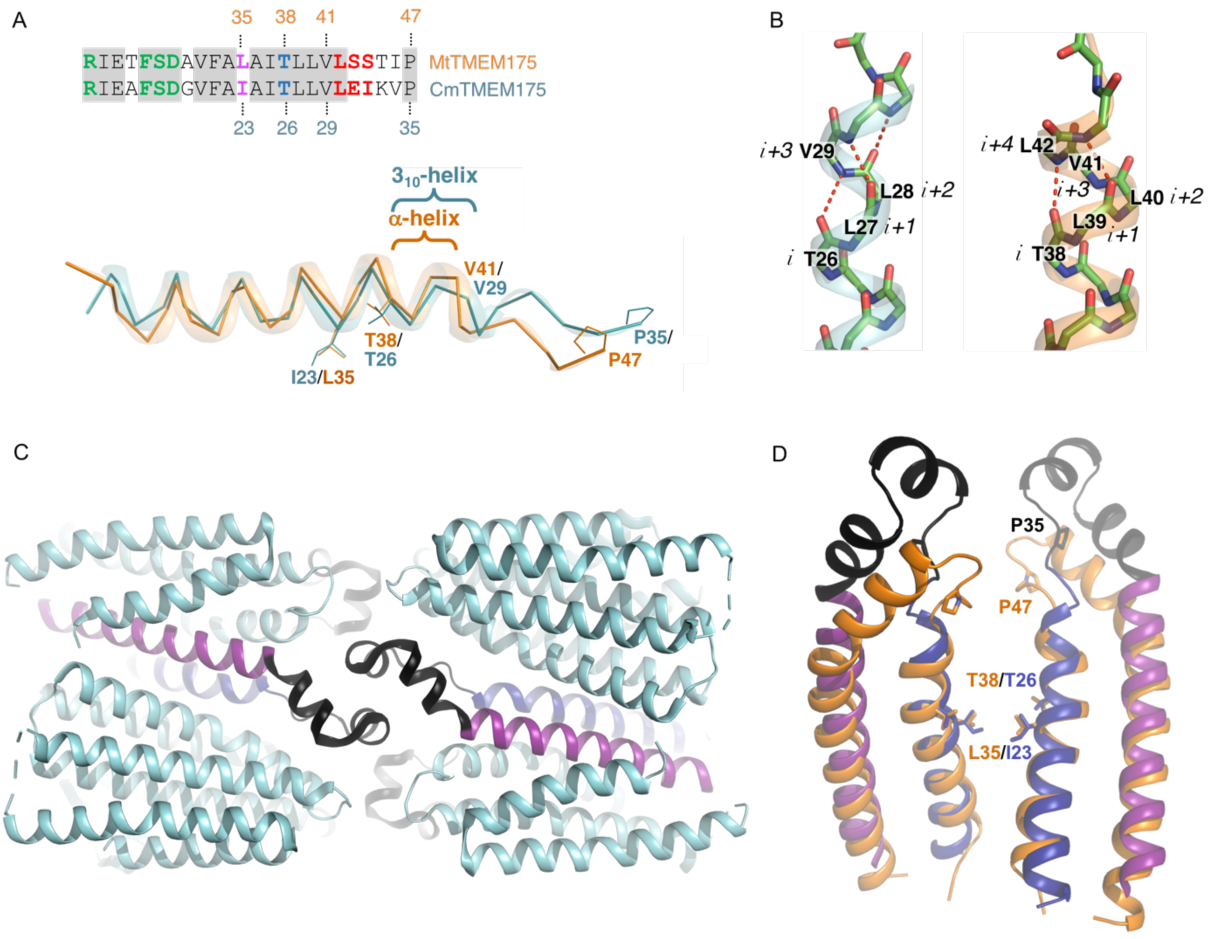


**Fig. S12. $3_{10}$-helix and crystal contacts in the structure of CmTMEM175. (A)** Overlay of helix 1 in MtTMEM175 (orange) and CmTMEM175 (cyan, PDB accession 5VRE) in ribbon representation. The structure of CmTMEM175 features a $3_{10}$-helix at the extracellular end whereas the corresponding region in MtTMEM175 is alpha helical. Numbering of selected residues is indicated. A sequence alignment of the respective region is shown. Coloring in the alignment is as in figure S9B. **(B)** Extracellular tips of helix 1 in CmTMEM175 (cyan) and MtTMEM175 (orange) shown as stick/ cartoon representation. In CmTMEM175 three residues complete a helical turn ($3_{10}$-helix) while in MtTMEM175 four residues complete a helical turn (alpha helical). Only backbone atoms are displayed and involved residues are indicated. **(C)** Crystal contacts in the CmTMEM175 structure. Helix 1 is colored in dark blue, helix 2 in magenta and the loop connecting helix 1 and 2 participating in crystal contacts is depicted in black. **(D)** Overlay of helix 1, 2 and connecting loops of CmTMEM175 (same colors as in **(C)**) and MtTMEM175 (orange). Selected residues in helix 1 are shown.

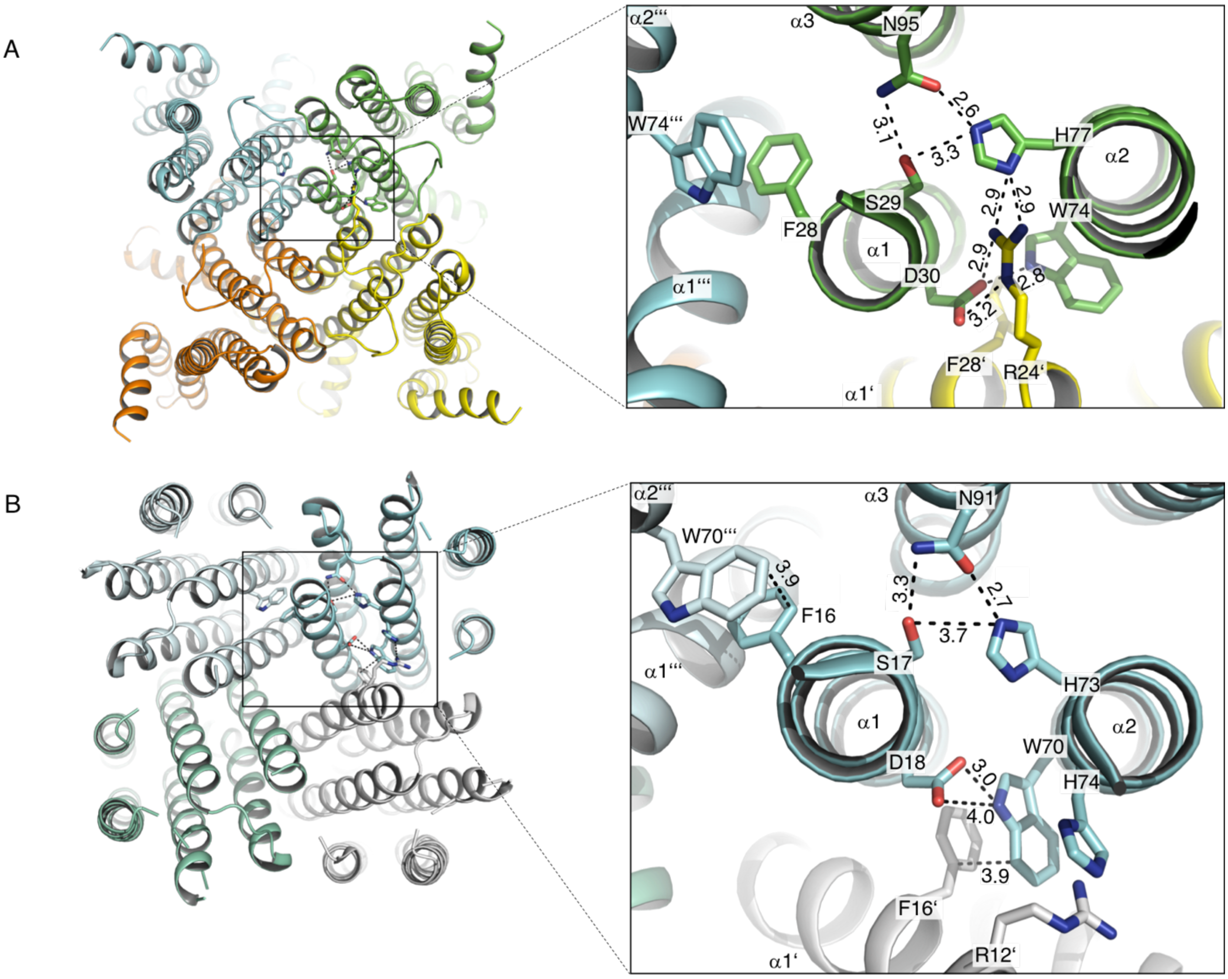


**Fig. S13. Interactions between conserved residues in TMEM175 proteins. (A)** MtTMEM175. **(B)** CmTMEM175 (5VRE). Interacting residues are shown with corresponding numbering and distances are given in Å. Individual subunits are emphasized with different colors. The view is from intracellular.

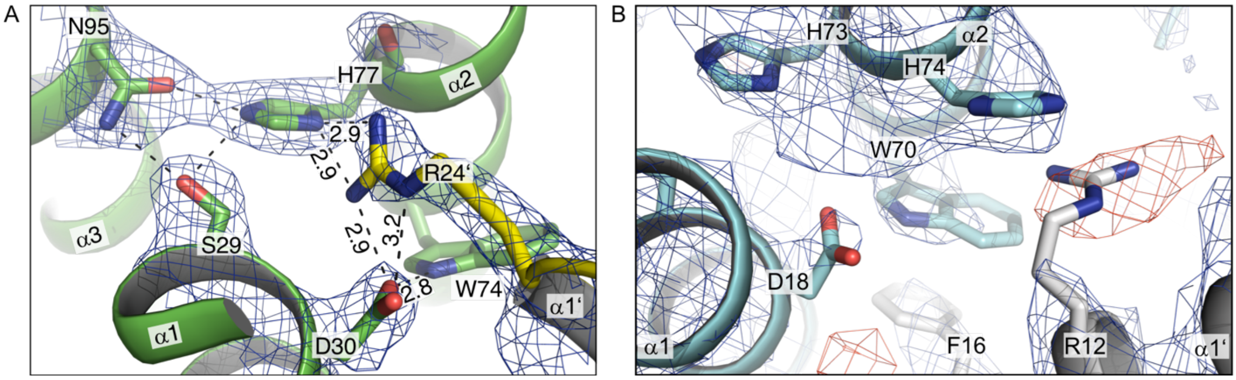


**Fig. S14. Interactions of the conserved arginine at the N-terminus of helix 1. (A)** Close-up view of $Arg^{24}$ in helix 1 of MtTMEM175 from intracellular, showing the interaction with $His^{77}$ and $Asp^{30}$ of the adjacent subunit. Bond distances are indicated in Å. The $2F_o$-$F_c$ electron density is shown as blue mesh (at 2.4 Å, contoured at 1.8 σ, sharpened with b=-25). **(B)** Negative difference electron density in the structure of CmTMEM175 (5VRE) at the position of $Arg^{12}$. The $2F_o$-$F_c$ electron density (at 3.3 Å, contoured at 1.55 σ, blue) and the $F_o$-$F_c$ density (contoured at -3 σ, red) are depicted. The view is from intracellular.

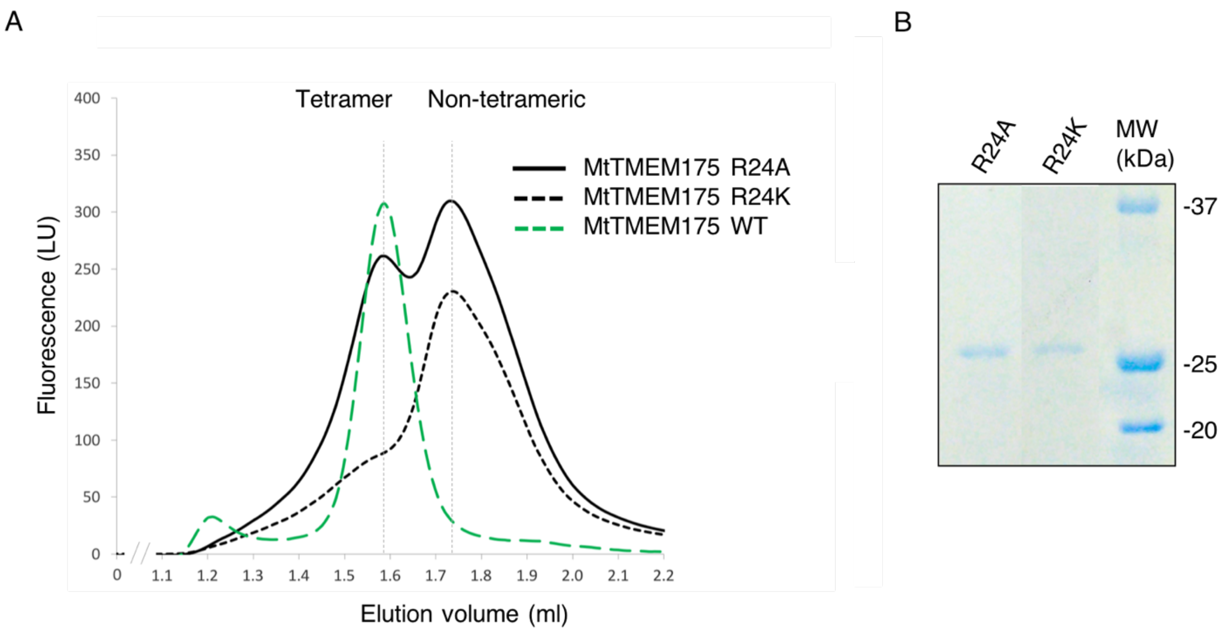


**Fig. S15. Defective tetrameric assembly in mutants of Arg$^{24}$ in MtTMEM175. (A)** Size exclusion chromatograms from a Superdex 200 5/150 column of MtTMEM175 Arg$^{24}$ mutant proteins compared to a WT chromatogram. The peak height of WT MtTMEM175 is normalized to the peak height of the mutant R24A. **(B)** Coomassie-stained SDS-PAGE gel of purified R24A and R24K mutant proteins that were subjected to SEC in **(A)**. The molecular weight marker (MW) is shown on the right.

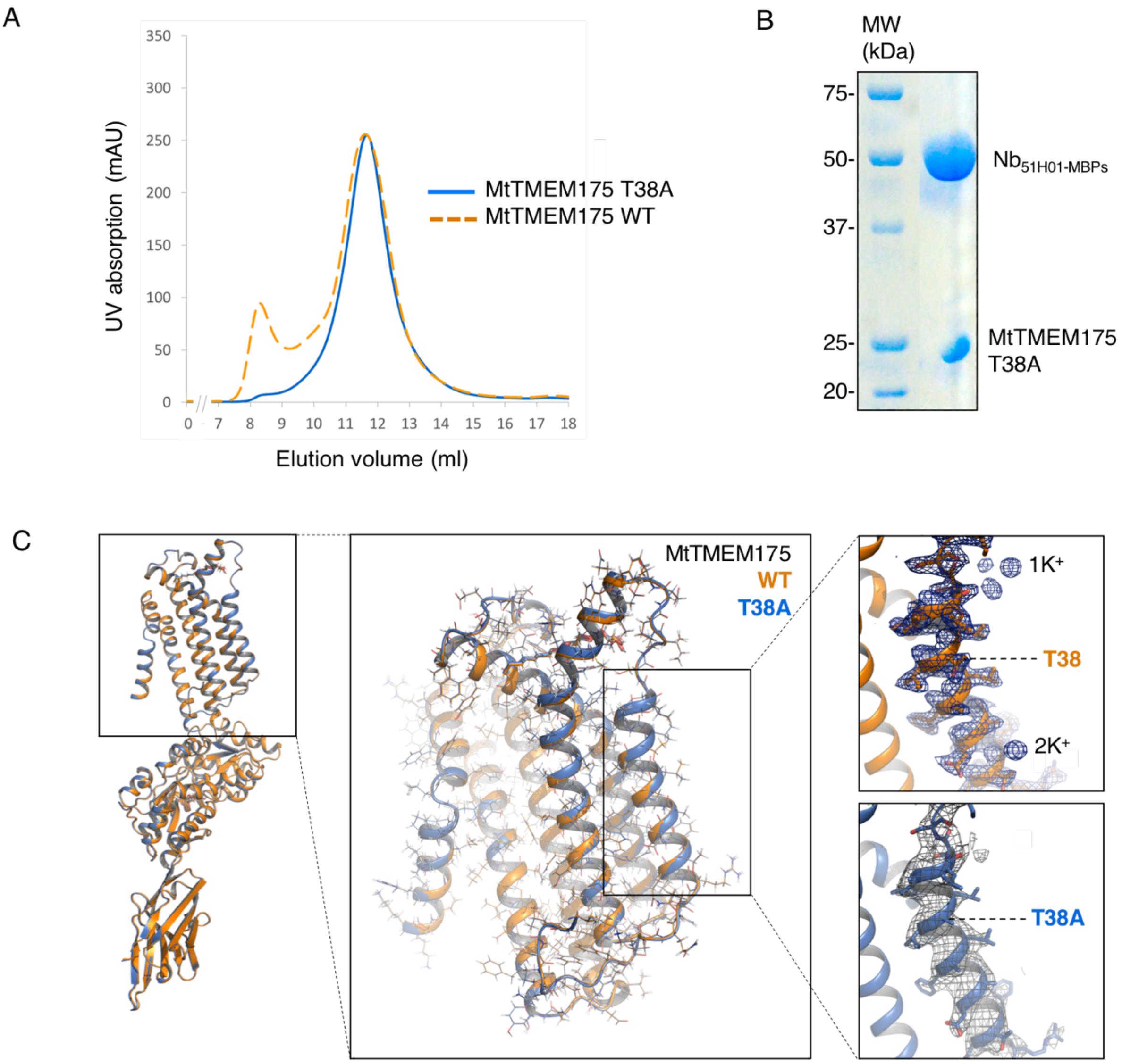


**Fig. S16. Crystallization of the T38A mutant of MtTMEM175. (A)** Size exclusion chromatogram of MtTMEM175 carrying a T38A substitution and WT for comparison (left). The peak height of the WT is normalized to the peak height of the mutant. **(B)** Coomassie-stained SDS-PAGE gel of the MtTMEM175-$Nb_{51H01\text{-}MBPs}$ complex after size exclusion chromatography before setting up crystals. The molecular weight marker (MW) is shown on the left. **(C)** Overlay of mutant and WT MtTMEM175 structures and close-up view on helix 1. Only one subunit is shown. Key residues $Thr^{38}$/ T38A are indicated. WT in orange, and mutant in blue. The $2F_O$-$F_C$ density of WT (at 2.4 Å, contoured at 1.8 σ after sharpening with b=-25, blue) and of the T38A mutant (at 3.4 Å, contoured at 1.3 σ, grey) are shown (see also Table S2, T38A).

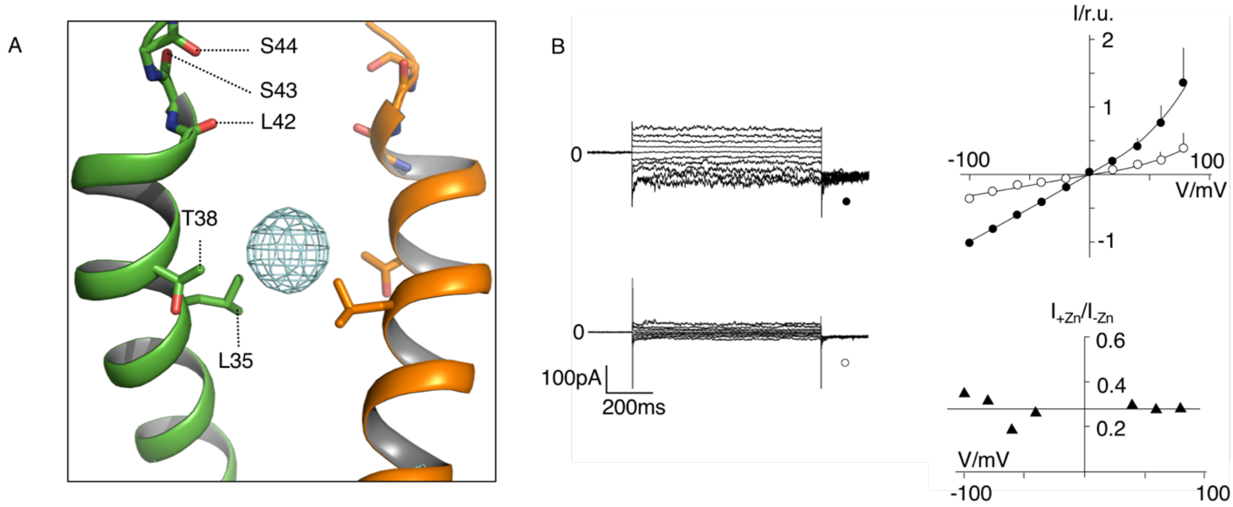


**Fig. S17. Anomalous density of zinc in WT MtTMEM175 and zinc block in the mutant T38A. (A)** Location of zinc ions within the pore. Anomalous difference electron density of $Zn^{2+}$ is illustrated as cyan mesh (at 2.88 Å, contoured at 4 σ, blurred with b=170). Front and rear subunits are omitted for clarity. **(B)** Currents recorded in HEK293 cells expressing the MtTMEM175 T38A mutant (left row) before (top) and 5 min after adding 5 mM $ZnSO_4$ (bottom) to the bath solution with 150 mM $K^+$. Right panel shows mean I/V relation (top) of n=3 cells (± s.d.) before (●) and after (○) adding 5 mM $ZnSO_4$ to the bath solution (top). To compare the effect on different cells the I/V relation was normalized to currents at -100 mV in the absence of blocker. The ratio of currents without and with blocker ($I_{-Zn}/I_{+Zn}$) is voltage independent (bottom).

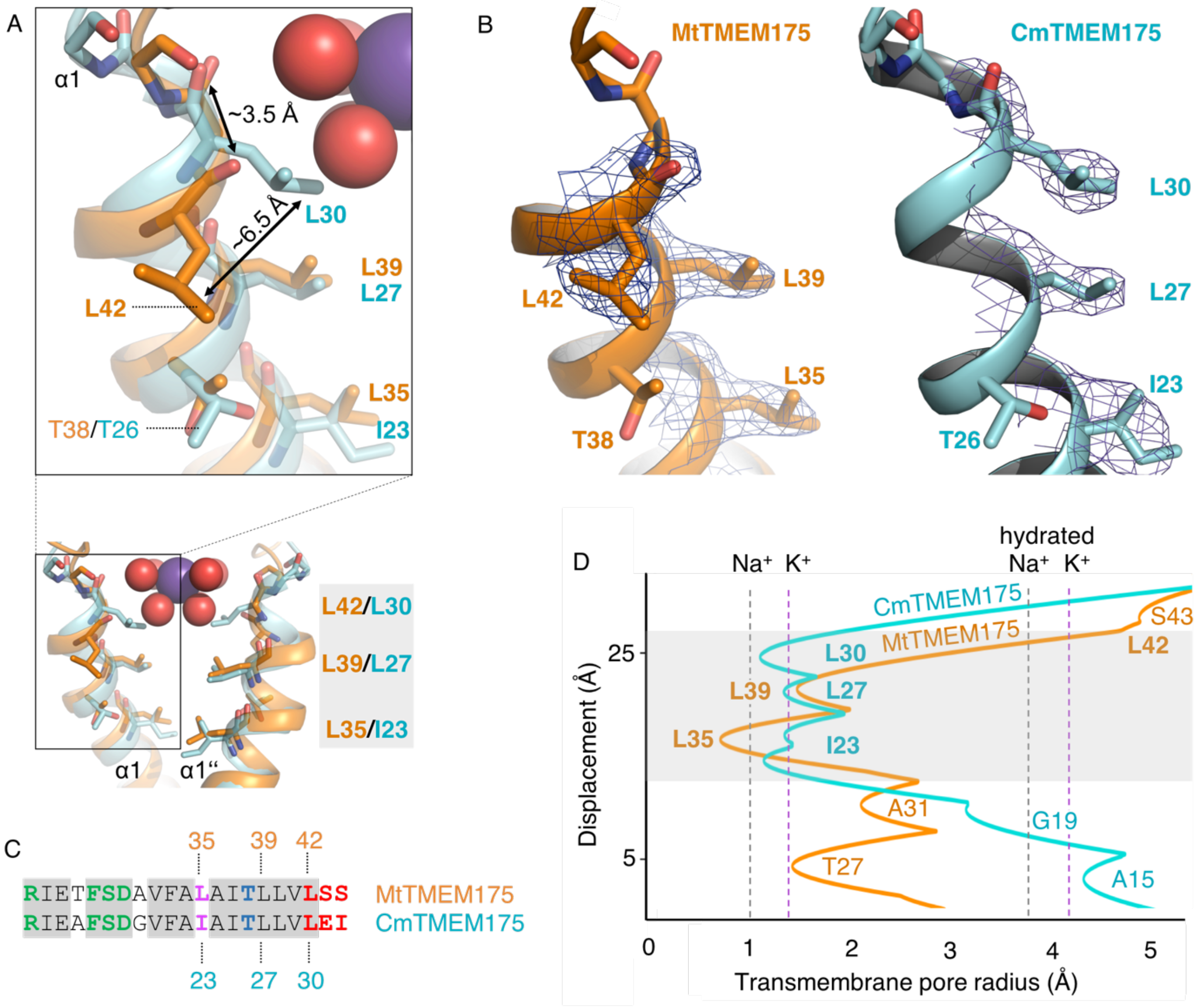


**Fig. S18. Constriction points in MtTMEM175 and CmTMEM175. (A)** Overlay of helix 1 from MtTMEM175 (orange) and CmTMEM175 (5VRE, cyan) with close-up view on residues Leu$^{35}$, Leu$^{39}$ and Leu$^{42}$ in MtTMEM175 and Ile$^{23}$, Leu$^{27}$ and Leu$^{30}$ in CmTMEM175, respectively. Deviations between the side chains and backbone oxygens of Leu$^{42}$ in MtTMEM175 and Leu$^{30}$ in CmTMEM175 are shown in Å. The view is from the side. In **(B)** the corresponding $2F_o$-$F_c$ densities are shown for MtTMEM175 (left, at 2.4 Å, contoured at 1.8 σ after sharpening with b=-25) and CmTMEM175 (right, at 3.3 Å, contoured at 1.55 σ). **(C)** Sequence alignment of helix 1 in MtTMEM175 and CmTMEM175 with respective residues numbered. Coloring is as in figure S9B. **(D)** Comparison of a HOLE analysis of the pore in MtTMEM175 and CmTMEM175. The pore radius along the central axis is shown in Å. Dashed lines indicate the radii for $K^+$ and $Na^+$ with and without hydration shell. The structures used in the HOLE analysis were aligned to superimpose Leu$^{35}$ and Thr$^{38}$ in MtTMEM175 with Ile$^{23}$ and Thr$^{26}$ in CmTMEM175.

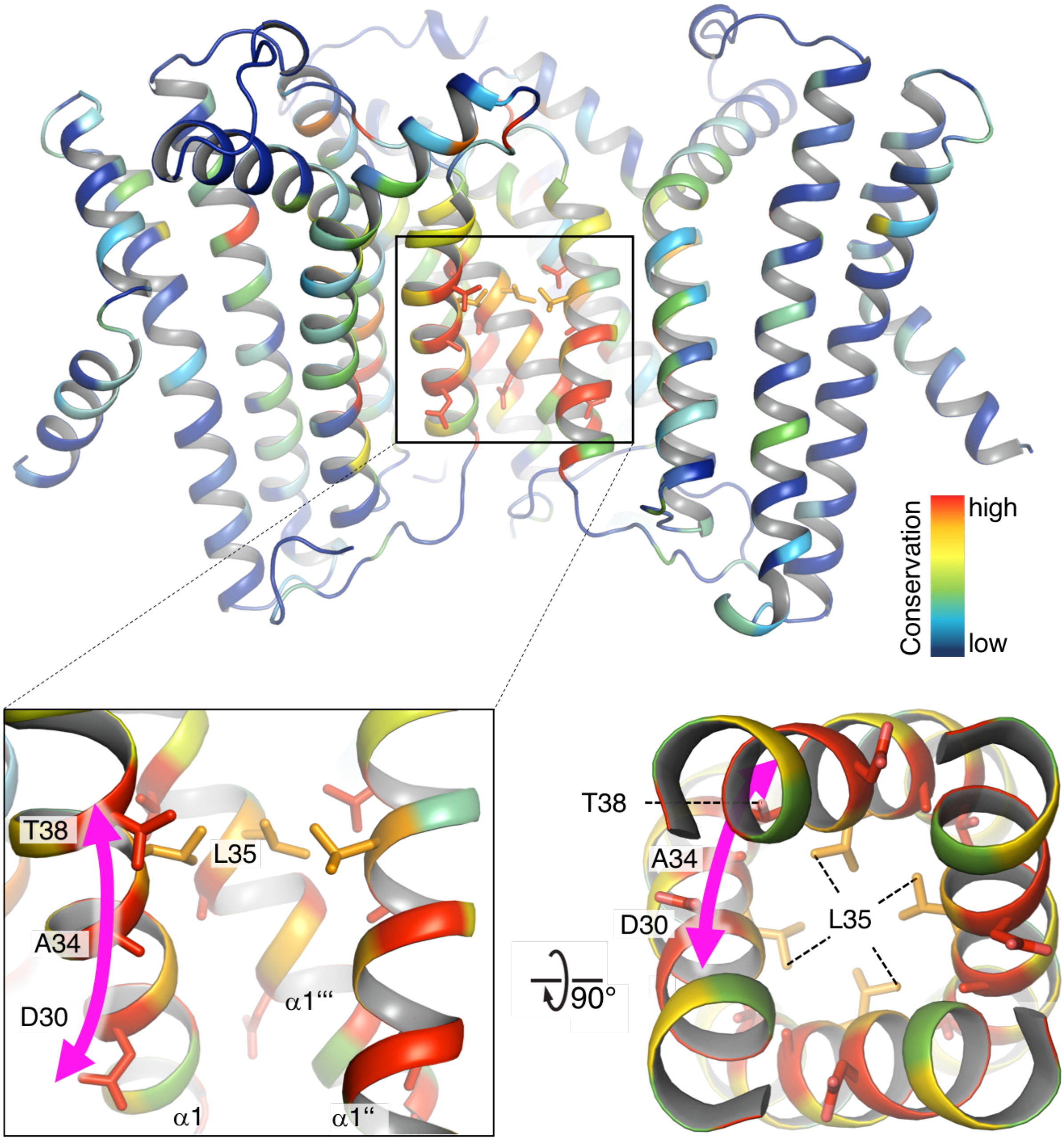


**Fig. S19. Conservation in TMEM175 projected onto the structure of MtTMEM175**. Top: Side view of the channel is shown with enlarged view (boxed, bottom left). One subunit has been omitted for clarity. Bottom right: View from intracellular on the pore. Conservation is color-coded in rainbow colors. Pink arrows mark the course of small highly conserved residues on helix 1 ($Asp^{30}$, $Ala^{34}$, $Thr^{38}$) (see also fig. S9 and Supplemental Material and Methods). Pore-lining $Leu^{35}$ is shown for orientation.

| | Native | K/S | Cs | Rb | Zn |
|---|---|---|---|---|---|
| **PDB Code** | **XXXX** | **XXXX** | **XXXX** | **XXXX** | **XXXX** |
| **Data collection** | | | | | |
| Wavelength (Å) | 0.87933 | 2.02460 | 2.00690 | 0.79480 | 1.24610 |
| Space group | $P4\,2_1\,2$ | $P4\,2_1\,2$ | $P4\,2_1\,2$ | $P4\,2_1\,2$ | $P4\,2_1\,2$ |
| Unit cell dimensions | | | | | |
| a, b, c (Å) | 131.2 131.2 132.6 | 132.0 132.0 132.6 | 131.7 131.7 131.5 | 130.5 130.5 132.4 | 131.9 131.9 132.7 |
| α, ß, λ (°) | 90, 90, 90 | 90, 90, 90 | 90, 90, 90 | 90, 90, 90 | 90, 90, 90 |
| Resolution (Å) | 46.6-2.4 (2.49-2.4)* | 46.8-3.5 (3.63-3.5)* | 46.5-3.8 (3.94-3.8)* | 46.5-3.5 (3.62-3.5)* | 46.64-2.90 (3.0-2.9)* |
| $R_{meas}$ (%) | 29.1(415.3) | 6.8 (128.3) | 24.8 (776.8) | 27.9 (351.2) | 17.7(1,432) |
| $R_{pim}$ (%) | 2.2 (43.1) | 1.8 (30.3) | 1.9 (41.7) | 3.3 (74.0) | 2.6 (223.0) |
| I/σI | 22.7(1.5) | 15.6 (2.0) | 10.2 (1.0) | 11.4 (1.4) | 10.2 (0.60) |
| $CC_{1/2}$ | 100.0 (49.4) | 99.6 (88.6) | 100.0 (75.7) | 100.0 (51.1) | 99.9 (46.7) |
| Completeness (%) | 99.3 (99.4) | 99.8 (98.3) | 99.9 (100.0) | 99.3 (93.2) | 99.9 (99.8) |
| Redundancy | 166.6(83.2) | 9.6 (9.3) | 40. (40.0) | 27.7 (23.0) | 27.0 (27.1) |
| Unique reflections | 45,710 (4500) | 28,188 (2080) | 11,875 (1153) | 14,836 (1400) | 26,575 (2,588) |
| **Refinement** | | | | | |
| Resolution (Å) | 24.16-2.40 | | 30.21-3.80 | 21.47-3.50 | 44.1-2.90 |
| $R_{work}/R_{free}$ | 0.209/0.253 | | 0.280/0.337 | 0.289/0.291 | 0.275/0.286 |
| Ramachandran favored/outlier | 95.9/0.0 | | 94.0/0.3 | 95.9/0.0 | 95.3/0.2 |
| No. atoms | 5,670 | | 5,549 | 5,532 | 5,548 |
| Protein | 5,463 | | 5,463 | 5,463 | 5,463 |
| Ligand/ion | 55 | | 54 | 54 | 56 |
| Water | 152 | | 32 | 15 | 29 |
| B-factors | 132.7 | | 170.2 | 173.3 | 161.8 |
| Protein | 134.1 | | 171.0 | 174.2 | 162.2 |
| Ligand/ion | 91.7 | | 93.5 | 93.7 | 141.5 |
| Water | 95.1 | | 158.7 | 156.0 | 124.6 |
| R.m.s. deviations | | | | | |
| Bond lengths (Å) | 0.014 | | 0.013 | 0.016 | 0.013 |
| Bond angles (°) | 1.57 | | 1.49 | 1.82 | 1.52 |
| Clashcore | 0.54 | | 1.36 | 1.27 | 0.09 |

*Values in parentheses are for highest-resolution shell.

**Table S1. Crystallographic data collection and refinement statistics.** The resolution cutoff was determined by $CC_{1/2}$ criterion (*57*). For the native data set seven datasets from a single crystal were merged together. For the K/S data set, two sets from a single crystal have been merged together. For the Cs dataset, three datasets from three crystals have been merged together. Six data sets from two crystals have been merged together for the Rb data set. For the Zn data set, two data sets from two crystals have been merged together.

| | **T38A** |
|---|---|
| **PDB Code** | **XXXX** |
| **Data collection** | |
| Wavelength (Å) | 1.00004 |
| Space group | $P4\ 2_1\ 2$ |
| Unit cell dimensions | |
| a, b, c (Å) | 133.5 133.5 132.5 |
| α, ß, λ (°) | 90, 90, 90 |
| Resolution (Å) | 47.2-3.40 (3.67-3.40)* |
| $R_{meas}$ (%) | 255.2 (10,621) |
| $R_{pim}$ (%) | 21.6 (901.5) |
| I/σI | 6.5 (1.4) |
| $CC_{1/2}$ | 99.9 (79.7) |
| Completeness (%) | 99.9 (99.6) |
| Redundancy | 146.0 (144.6) |
| Unique reflections | 16,953 (3435) |
| **Refinement** | |
| Resolution (Å) | 22.25-3.40 |
| $R_{work}/R_{free}$ | 0.273/0.292 |
| Ramachandran favored/outlier | 95.9/0.0 |
| No. atoms | 5,542 |
| Protein | 5,461 |
| Ligand/ion | 54 |
| Water | 27 |
| B-factors | 178.9 |
| Protein | 179.6 |
| Ligand/ion | 150.7 |
| Water | 101.8 |
| R.m.s. deviations | |
| Bond lengths (Å) | 0.016 |
| Bond angles (°) | 1.81 |
| Clashcore | 0.99 |

*Values in parentheses are for highest-resolution shell.

**Table S2. Crystallographic data collection and refinement statistics (T38A).** The resolution cutoff was determined by $CC_{1/2}$ criterion (*57*). For the T38A mutant dataset, six datasets from two crystals have been merged together.
